# Vascular Smooth Muscle Myosin 2 Filaments Dynamically Assemble and Stabilize During Induced Contractility

**DOI:** 10.1101/2022.10.10.511341

**Authors:** Sasha K. Demeulenaere, Margaret A. Bennett, Bradley Somerfield, Huini Wu, Margaret E. Utgaard, Hiral Patel, Stefano Sala, Elizabeth R. Longtine, Ahmed Zied, Jonathan A. Kirk, Patrick W. Oakes, Jordan R. Beach

## Abstract

**Background:** Vascular smooth muscle cells (SMCs) dynamically tune blood vessel diameter to regulate blood pressure, provide vessel wall structural integrity, and absorb shock on a beat-to-beat timescale. Smooth muscle myosin 2 (SMII) is the dominant motor protein driving SMC contraction. To function, SMII monomers dynamically assemble into filaments, which associate with the actin cytoskeleton to drive contractility. Precisely how SMII filaments assemble and exchange in living SMCs, however, both at steady-state and during induced contractility, remains poorly defined.

**Methods:** We used a single-cell filament assembly assay to determine SMII assembly into filaments at steady-state and upon induced contractility in rat aortic SMCs (A7R5) transiently-expressing EGFP-tagged SM1A isoform of SMII. We then used fluorescence recovery after photobleaching (FRAP) to characterize SMII exchange kinetics at steady-state and upon induced contractility, and measured changes in force production using traction force microscopy. Finally, we developed a CRISPR knock-in EGFP-SMII murine model to quantify SMII dynamics at endogenous expression in primary SMCs and intact arterioles.

**Results:** While predominantly filamentous at baseline, induced contraction rapidly increased SMII filament assembly. FRAP revealed rapid SMII exchange kinetics, more similar to non-muscle myosin II than striated myosin II, and induced contractility consistently stabilized SMII filaments. Super-resolution imaging revealed SMII and non-muscle myosin II filament structures consistent with co-assembly. Endogenous EGFP-SMII in primary SMCs and intact arterioles paralleled cell culture studies with similar baseline exchange kinetics and activation-dependent stabilization.

**Conclusions:** Together, these data support a model in which SMII is surprisingly dynamic and co-assembles with non-muscle myosin II. Vascular SMC activation further increases SMII filament assembly while reducing filament exchange, consistent with stabilization of a dynamic SMII pool during force generation, which allows cells to dynamically adapt their overall contractility in response to environmental conditions.

**CLINICAL IMPLICATIONS:** Smooth muscle myosin II (SMII) is the principal contractile motor protein of vascular smooth muscle tissue, but its molecular behavior in living differentiated SMCs remains poorly defined. Here, we show that SMII is not a static contractile scaffold but highly dynamic. SMII exhibits dynamic exchange between the filament and monomeric forms at baseline, but becomes acutely stabilized during agonist-induced contraction, including in primary SMCs and intact arterioles at endogenous *Myh11* expression. These findings suggest that vascular tone is tuned not only by activating pre-existing myosin filaments, but also by rapidly shifting SMII filament assembly levels and exchange rates. This framework is relevant to diseases in which SMC contractility and arterial wall mechanics are abnormal, including *MYH11*-associated thoracic aortic disease and related disorders of arterial stiffness or maladaptive vasomotor regulation. Although this study is mechanistic, it identifies SMII filament dynamics as a measurable layer of vascular regulation and a potential readout for future studies of pathogenic *MYH11* variants, disease modeling, and therapies that alter myosin activation and/or cytoskeletal stability.

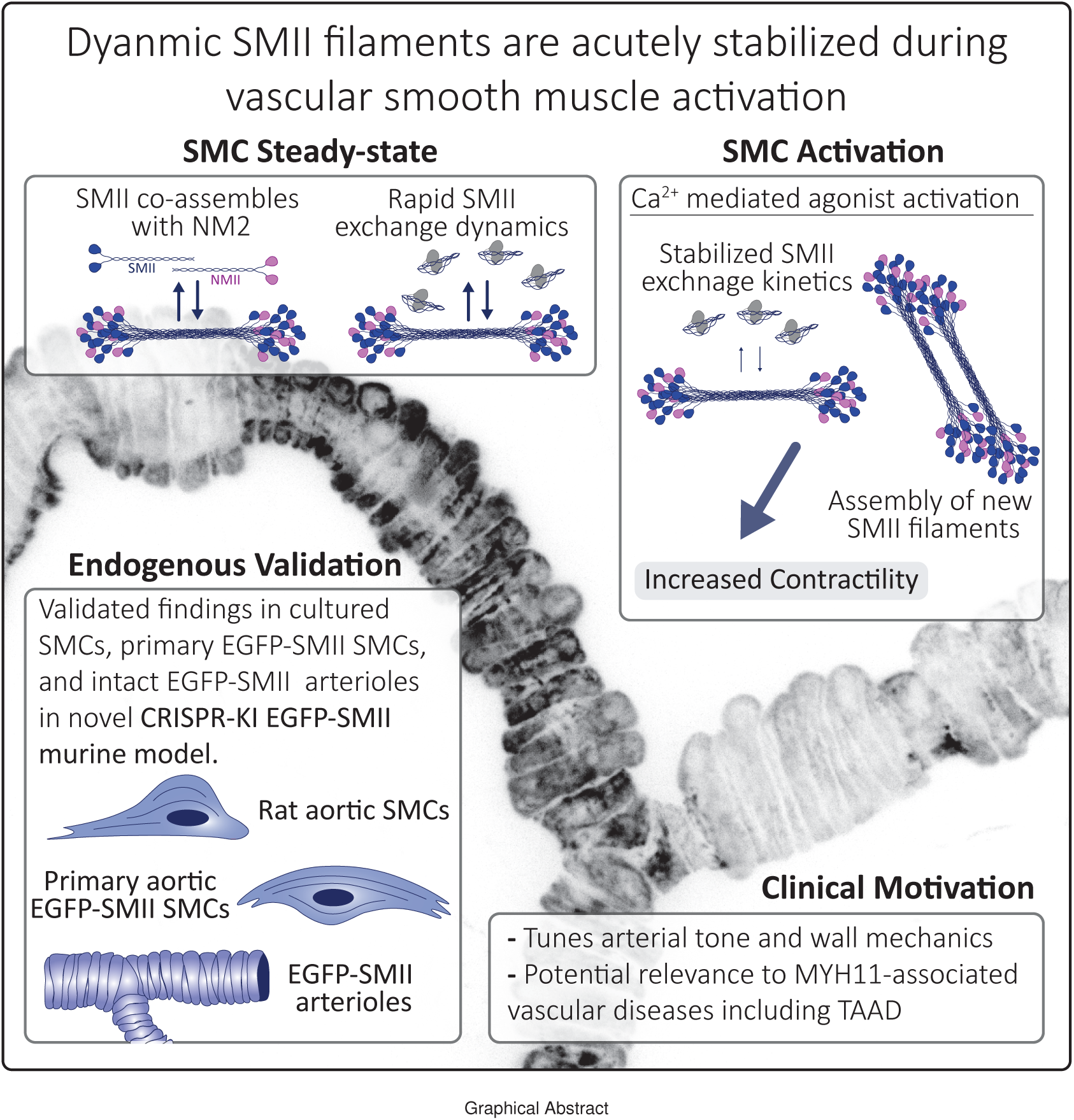

## INTRODUCTION

Smooth muscle is a fundamental component of the cardiovascular system. In arteries, the smooth muscle layer (tunica media) comprises a significant portion of the vessel wall and is dominated by vascular smooth muscle cells (SMCs). SMCs provide structural integrity and dynamically contract and relax to alter vessel diameter, thereby regulating vascular tone and blood pressure and absorbing pulsatile mechanical loads on a beat-to-beat timescale (1). Contraction in SMCs is primarily powered by the dominant contractile protein smooth muscle myosin 2 (SMII) acting on the actin cytoskeleton (2). Consistent with SMII’s central role in SMC contractile function, rare variants in SMII and other contractile proteins are linked to inherited aortic disease, including aneurysms and dissections (3–5). SMC function, therefore, is critical to vascular pathophysiology. How SMII filaments dynamically assemble and exchange in living SMCs, however, remains poorly defined. Filling this knowledge gap is essential for understanding how SMII supports physiological contraction and how SMII disruption contributes to disease.

SMII is a member of the myosin 2 family, which also consists of striated myosin 2s (skeletal and cardiac) and non-muscle myosin 2s (NM2) (6). All myosin 2 “monomers” consist of two dimerizing heavy chains, two essential light chains, and two regulatory light chains (RLC). The extended coiled-coil of the heavy chain can reversibly associate with other monomers in parallel and antiparallel fashion to enable the assembly of bipolar filaments with motor domains at opposing ends (Fig. 1A; (7, 8)). These filaments are the functional units that interact with filamentous actin to drive contractile events. In SMCs, contractile activation is classically controlled by calcium-dependent and RhoA/ROCK-dependent pathways that regulate myosin RLC phosphorylation (9–11). Because SMII filament formation is reversible, the kinetics of exchange between monomer and filament-associated SMII pools can tune contractility on acute timescales.

**Fig. 1.**
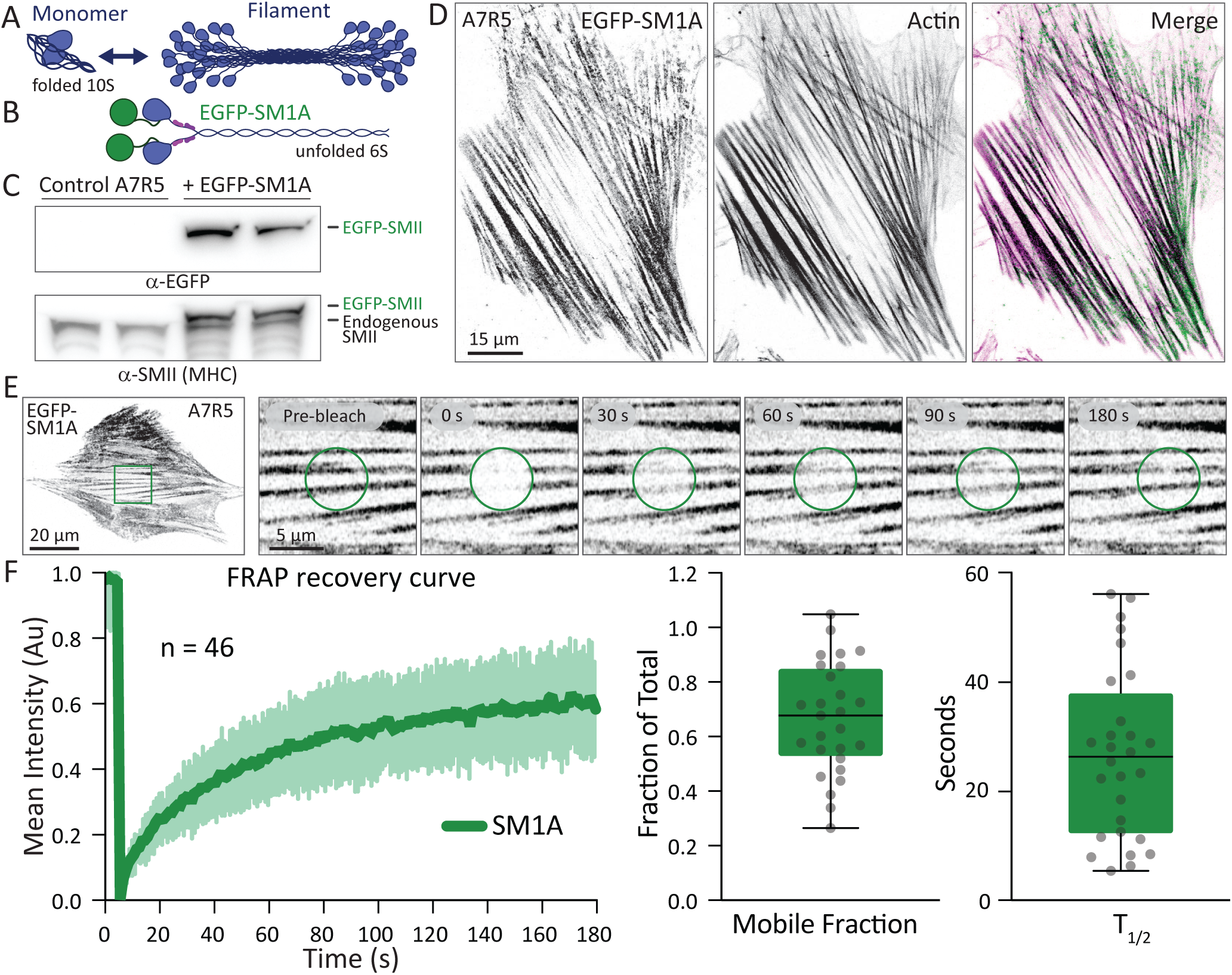
EGFP-SM1A filaments in A7R5 are highly dynamic. **A**) Diagram of myosin 2 in folded monomeric and filamentous structural states. **B)** Diagram of an SMII unfolded monomer tagged on the N-terminus with EGFP. **C)** Duplicate western blot of untransfected A7R5 (left lanes) and transiently expressing EGFP-SM1A (right lanes) probed with *α*-EGFP or *α*-SMII. **D)** Example images of an A7R5 cell transiently expressing EGFP-SM1A (green), fixed and stained with phalloidin (actin; magenta). **E)** FRAP example of an A7R5 cell expressing EGFP-SM1A. Bleach region indicated by green circle in insets. **F)** Average FRAP recovery curve plotted for EGFP-SM1A from 3 independent experiments, along with mobile fraction and t^1^⁄2 data displayed as box and whiskers (median *±* quartiles), calculated using a first-order exponential fit.

Myosin 2 family members appear to modulate assembly dynamics and total assembly levels to differing extents. Here, we use the term “assembly level” to refer to the fraction of cellular SMII residing in filamentous structures versus the mobile monomeric pool. For example, non-muscle cells rapidly modulate NM2 filament assembly while maintaining about half of the total myosin 2 pool in filamentous form (12, 13). In contrast, striated myosin 2s assemble more stable filaments that generate contractile force independent of new filament assembly, with the vast majority of the total myosin 2 pool in filamentous form (14). While significant contributions have been made to the field in recent decades(15, 16), we have less mechanistic insight into the molecular details of SMII filament assembly dynamics and activation than we do for striated and non-muscle myosin 2s. It has been suggested that SMC contraction primarily reflects activation of preassembled SMII filaments (17), and that assembly in tissue can be stimulated by changes in muscle length (18). At the cellular level, SMII exists in both monomer and filament form (19), however, and it is unclear whether SMII filaments in SMCs are stable (striated-like) or exchange dynamically (NM2-like), and how agonist activation modulates assembly and exchange. Considering SMII is genetically similar to NMII, but perhaps more functionally similar to striated myosin 2s, which are stably assembled filaments to drive repeated uniaxial contractile events, SMII behavior in cells is difficult to predict without experimentation.

A defining feature of vascular SMCs is their phenotypic plasticity, which enables adaptive tissue remodeling and repair in response to changing environmental cues. Vascular SMCs exist on a functional spectrum between a differentiated “contractile” phenotype and a more dedifferentiated migratory “synthetic” phenotype (20, 21). The differentiated phenotype is relatively stable and described as quiescent, whereas the dedifferentiated phenotype is characterized by increased proliferative and migratory ability. Previous studies have shown that SMCs can dedifferentiate rapidly in tissue culture (22, 23). Because phenotype transitions can alter the expression, localization, and regulation of contractile proteins, measurements in standard *in vitro* models may not fully reflect SMII behavior in differentiated SMCs *in vivo*. Accordingly, investigating endogenous SMII in primary cells and intact tissue is ideal to define physiologic assembly and exchange dynamics in their native context.

In this work, we first use exogenous expression of the dominant vascular SMII isoform (SM1A) in cell culture models to demonstrate that SMII is highly dynamic but primarily filamentous at steady-state, and that agonist-induced SMC contraction leads to a hyper-assembled but more stable SMII network. This response is paralleled with cytoplasmic calcium release and cell-scale force production. High resolution imaging additionally suggests co-assembly with NM2. Complementary and consistent results were obtained with endogenous SMII in primary SMCs and intact arterioles using a novel EGFP-SMII knock-in murine model. Together, our data support a model in which SMC activation induces both nascent SMII filament assembly and activation of pre-existing filaments.

## RESULTS

### SM1A and NM2 filaments in cultured SMCs have dynamic exchange kinetics and likely co-assemble

A single gene (*MYH11*) produces four dominant smooth muscle splice variants (SM1A, SM1B, SM2A, SM2B) that vary in the presence or absence of splice inserts in the motor domain or C-terminal tail (24). The dominant isoform in vascular SMCs is SM1A (25, 26). Therefore, to better understand the assembly dynamics of SMII filaments in SMCs, we generated an expression plasmid with an EGFP coupled to the N-terminus of the SM1A heavy chain (EGFP-SM1A; Fig. 1B). Western blot analysis of transient overexpression of EGFP-SM1A in A7R5 demonstrated expression of the EGFP-SM1A shifted above endogenous SMII, as expected (Fig. 1C). High resolution imaging demonstrated that EGFP-SM1A associated with large actin stress fibers in cells (Fig. 1D). Therefore, our EGFP-SM1A plasmid appears to be a faithful reporter of SM1A in these immortalized SMCs. To investigate SMII filament exchange kinetics, we performed FRAP on A7R5 expressing EGFP-SM1A. Here, we use the term “exchange” to refer to the movement of myosin 2 monomers into and out of filaments, and avoid the term “turnover”, which might also refer to protein synthesis and degradation and should not be relevant during our experimental timescales of a few minutes. Fig. 1E shows an example FRAP experiment, with insets showing EGFP-SM1A before bleaching and during recovery. EGFP-SM1A displayed a relatively high fraction of the exchanging pool (mobile fraction ∼70%), and low time for half of the exchange to occur (t1⁄2 ∼ 30s) relative to striated myosin 2 paralogs (27, 28) (Fig. 1F). This suggests SM1A is capable of forming highly dynamic filaments in SMCs that are more reminiscent of non-muscle myosin systems (12, 13) than striated muscle myosin systems (27, 28).

### SMII co-assembles with NM2

As SMCs express both myosin 2 paralogs (29), we sought to carefully investigate their relationship in our cell model. We co-expressed EGFP-SM1A and NM2A-mApple in A7R5 cells and observed extensive overlap in filamentous structures throughout the cell (Fig. 2A). Closer examination of individual fibers revealed distinct alternating patterns of the two fluorophores (Fig. 2B). Because each paralog is tagged with their respective fluorophores at opposing ends of the monomer (N- and C-termini; Fig. 2B), this arrangement suggests that the SM1A and NM2A are co-assembling into filaments in the VSMCs, as has been seen previously for NM2 isoforms (30, 31). The alternating SMIA-NM2A-SM1A (green-magenta-green) patterns are also ∼300 nm in length (Fig. 2B), consistent with what would be seen in co-assembled filaments.

**Fig. 2.**
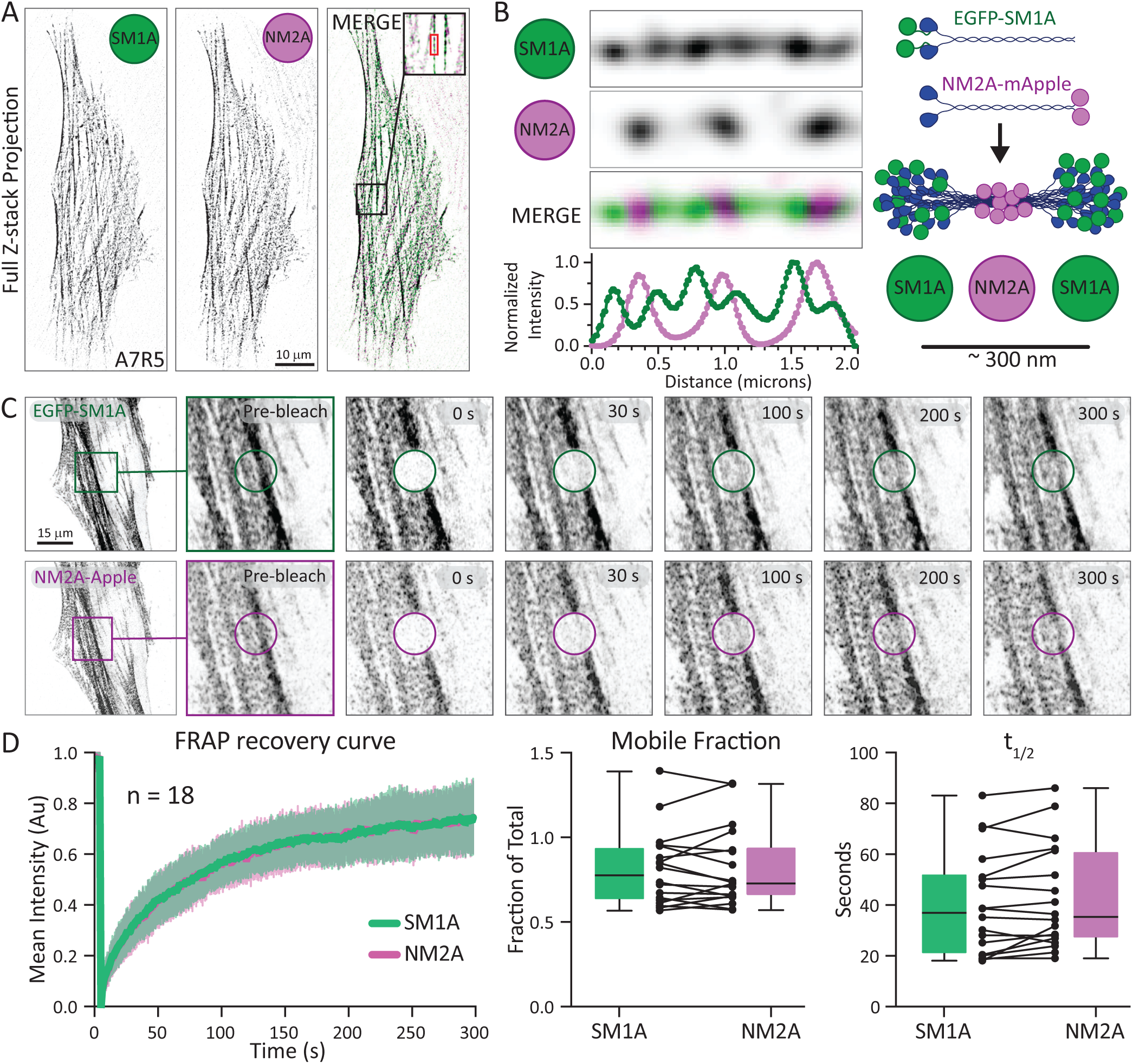
SM1A co-assembles with NMIIA. **A)** A7R5 cells co-transfected with EGFP-SM1A and NM2A-mApple were imaged with SIM and sum projected. Individual channels are displayed in inverted grayscale, and merge is shown in the right panel. **B)** An individual actomyosin fiber was cropped from the cell in (A) and displayed at higher zoom for SM1A, NM2 and merge. The lower plot is a line scan through the fiber normalized to maximum intensity for each channel. **C)** Diagram of co-assembled EGFP-SM1A and NM2A-mApple. D) FRAP example of an A7R5 cell expressing EGFP-SM1A (top row) and NM2A-mApple (bottom row). Bleach regions indicated by circles. Average recovery curves of SM1A and NM2 plotted as mean *±* SEM for 12 cells from 3 independent experiments. Mobile fraction and t^1^⁄2 are displayed as both box and whiskers (median *±* quartiles) and spaghetti plots. Dots in spaghetti plot represent individual cells for NM2 and SM1A with lines connecting cells. A paired t-test was performed and there was no statistically significant difference between groups.

We next asked if SM1A and NM2A in the same filamentous networks had similar exchange kinetics. We performed simultaneous FRAP in A7R5 cells expressing both EGFP-SM1A and NM2A-mApple (Fig. 2C). Our analysis revealed indistinguishable mobile fractions and t1⁄2 values (Fig. 2D), indicating that both SM1A and NM2A filaments exchange rapidly and readily when co-expressed in these cells. For SM1A, the exchange kinetics were similar regardless of NM2A expression (Figs. 1F, 2D), suggesting NM2 is not inducing a more dynamic SM1A. Collectively, then, SMII filaments in cultured SMCs display rapid polymer exchange kinetics that are indistinguishable from NM2 and co-localize with NM2 in filamentous structures in a manner consistent with co-assembly.

### A7R5 cells assemble new SMII filaments and display calcium-mediated contractile response to carbachol

Because A7R5 cells in culture are likely closer to a dedifferentiated SMC phenotype with reduced contractile capacity, we first sought to confirm that they could still increase their contractility in response to an agonist. We performed control experiments with the acetylcholine receptor agonist carbachol, a cholinergic receptor agonist that stimulates the release of intracellular calcium via IP3 (32). In GCaMP7s-expressing A7R5 cells, carbachol elicited transient increases in cytosolic calcium (Fig. 3A).

**Fig. 3.**
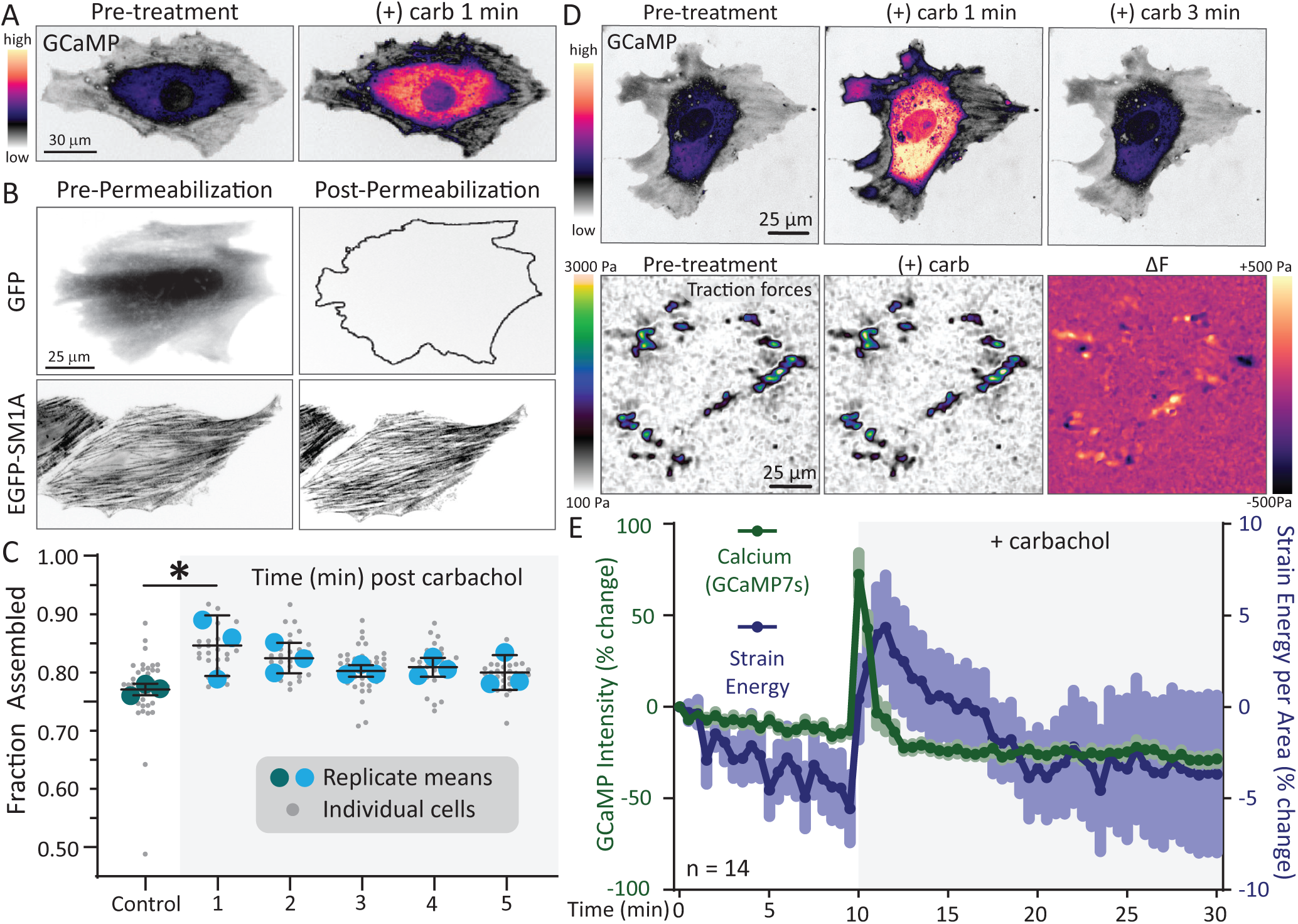
Induced contraction of SMCs drives SM1A assembly and a contractile response. **A)** A7R5 cell expressing GCaMP7s was imaged pre- and post-addition of 10 *μ*M carbachol. GCaMP intensity is displayed with iLUT (scale on left). **B)** A7R5 cells expressing EGFP-SM1A were imaged live to determine total SM1A intensity. Cells were treated with or without 10 *μ*M carbachol for the indicated time, monomeric SM1A was extracted using a triton buffer, and a second image was taken to determine the intensity of the remaining filamentous SM1A. As a control, A7R5 cells expressing free EGFP were imaged pre- and post-permeabilization to demonstrate cytoplasmic proteins are fully extracted from the cell. **C)** Quantification of the fraction of SM1A assembled in control cells and cells treated with 10 *μ*M of carbachol for the indicated time. Small circles indicate individual cells. Larger colored circles indicate means of three independent experiments. A one-way ANOVA was used with multiple comparisons between each time point and control to statistically analyze the data. * indidcates *p<* 0.05. **D)** An A7R5 cell expressing GCaMP7s was monitored for cytosolic calcium levels while simultaneously measuring its force production using traction force microscopy pre- and post-carbachol. Pre-carbachol and post-carbachol traction maps are the averages of the 10 min prior to and after carbachol treatment. The Δ*F* image represents post-carbachol - pre-carbachol average traction maps. **E)** Quantitation of the percent change of cytosolic calcium (green) and strain energy per area (blue) over time. Data plotted as mean *±* SEM for 15 cells from 3 experiments.

To determine if this calcium release produced SMII assembly, we performed a single-cell imaging-based triton permeabilization assay to measure populations of monomeric and filamentous SMII (33). A7R5 cells expressing EGFP-SM1A were first imaged to measure total EGFP-SM1A intensity. Then cells were treated with or without carbachol before permeabilizing with a physiological buffer containing triton to remove the soluble SMII pool. Finally, cells were re-imaged to measure remaining filamentous signal (Fig. 3B & C). Comparing intensity differences pre- and post-triton enables quantitative determination of monomer/filament ratios on a single-cell basis. Control experiments with A7R5 cells expressing free EGFP reveal a complete loss of EGFP signal upon triton permeabilization, demonstrating cytosolic proteins are completely liberated in this assay (Fig. 3B). Control assembly levels for SM1A at baseline were ∼75% filamentous (Fig. 3C& S1D). One minute after carbachol treatment, however, assembly levels increased by ∼10%, then decreased towards steady-state levels in the ensuing minutes (Fig. 3C). We did not observe any obvious changes in SMII-NM2 co-assembly upon carbachol treatment (Fig. S2), nor did observe any correlation between assembly and SMII intensity before permeabilization, suggesting the increase in assembly we observed with induced contractility was not affected by the expression level of exogenous EGFP-SMII. (Fig. S3C)

In parallel, we performed traction force microscopy (TFM) to determine if these cytosolic calcium and SMII assembly increases were harnessed into contractile energy. We observed contractile force generation following carbachol treatment with a slight delay and extended duration relative to the cytosolic calcium response (Fig. 3D&E), consistent with the calcium/MLCK-dependent signaling cascade that displays similar kinetics in intact smooth muscle tissue (34). Together, these data show that A7R5 cells mount a Ca^++^-dependent response to carbachol that is accompanied by a transient increase in SMII filament assembly and measurable contractile activity.

### SM1A filament dynamics are stabilized upon transiently-induced contraction

To determine if the increase in SMII assembly upon SMC induced contractility is accompanied by a change in filament exchange kinetics, we performed FRAP on a small region of an A7R5 expressing EGFP-SM1A, then treated with carbachol, and rapidly (within ∼1 min) performed FRAP on a similar but distinct region of the same cell (Fig. 4A). This provided cell-specific internal controls to monitor relative changes in assembly dynamics. Upon SMC activation, we observed a consistent decrease in mobile fraction, and a slight increase in t^1^⁄2, indicating a stabilization of the SMII machinery (Fig. 4B). To further validate our observed response to carbachol, we repeated the assay with an alternative agonist, angiotensin II (angII), which stimulates SMC contractility via angiotensin II receptor-mediated calcium release (35). We again observed a significant decrease in the mobile fraction, but with less impact on t^1^⁄2 (Fig. 4C). Together, internally-controlled FRAP measurements show that agonist stimulation (carbachol or angII) consistently reduces the SMII mobile fraction, indicating stabilization of the SMII filaments during induced contractility. **SMII exchange kinetics and assembly are unchanged upon differentiation of A7R5.** As SMCs are known to dedifferentiate in vitro, we sought to determine whether SM1A assembly dynamics were altered upon induction of the differentiated phenotype via serum starvation of A7R5s (36). Expression of contractile proteins in cells under various serum conditions shows that serum starvation increases both SMII and smooth muscle actin, confirming that cells were pushed towards the differentiated phenotype (Fig. S1A&B). FRAP analysis, however, revealed that SM1A exchange kinetics were not significantly altered, as there was no noticeable difference in mobile fraction or t^1^⁄2 between serum-starved or non-serum-starved cells (Fig. S1C). In our assembly assays, we also observed no significant difference in the steady-state assembled fraction between serum-starved and non-starved conditions (Fig. S1D). Together, these results suggest that while serum starvation increases expression of contractile markers and promotes a slightly more differentiated A7R5 phenotype, baseline SM1A filament assembly and exchange kinetics remain largely unchanged under these conditions.

**Fig. 4.**
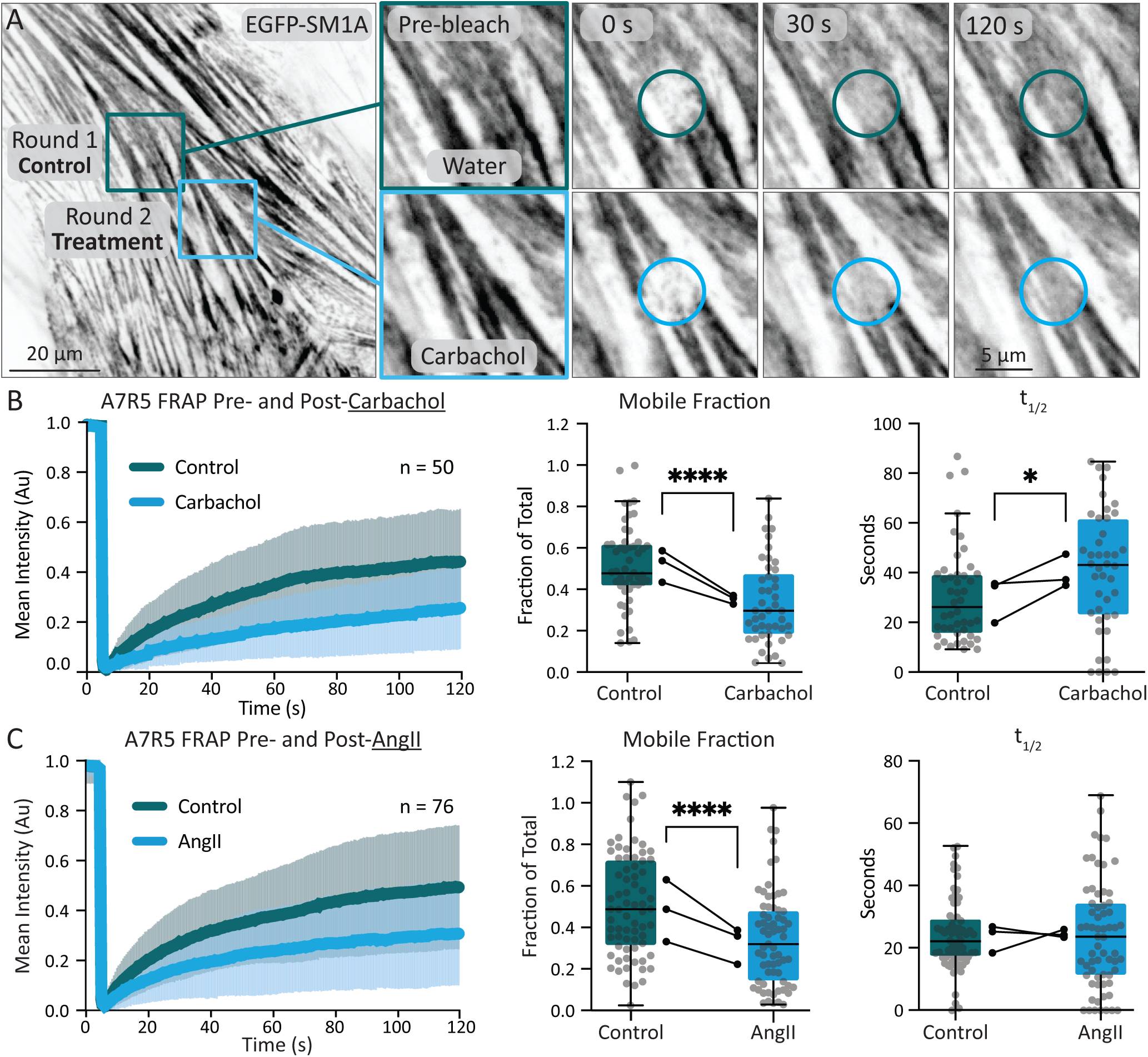
SM1A filament dynamics stabilize upon induced contraction. **A)** Example FRAP experiment of a single cell before and after addition of calcium agonist. Fluorescence recovery of EGFP-SM1A in A7R5 pre- (green) and post (blue) carbachol. Circles indicate the bleach region. **B)** Mean recovery curves from 50 individual cells across 3 independent experiments. Mobile fraction and t^1^⁄2 are reported with individual points in box and whiskers (median *±* quartiles). Means from each experiment are reported as dots in a spaghetti plot where pre- and post-carbachol measurements for individual experiments are connected. Paired t-tests were performed on control and drug treatment cells when measuring mobile fraction, t^1^⁄2. **C)** The same experiment as in (B) but using the agonist angII. Data represents 76 individual cells from 3 independent experiments. *, and **** represent *p<* 0.5 and *p<* 0.0001 respectively.

### Establishment of an EGFP-SMII murine model

While our data in A7R5 cells provide valuable insight into SMII assembly and exchange, SMC stable cell lines in culture are prone to phenotypic drift, and these measurements likely reflect SMII behavior in a more dedifferentiated state. To determine whether the same SMII organization and dynamics are preserved at endogenous expression in primary SMCs and intact tissue, we generated a knock-in mouse using CRISPR/Cas9 in which EGFP is fused in-frame to the N-terminus of the endogenous *Myh11* locus (Fig. 5A). As the splice variants do not use alternative translation start sites, this strategy labels all four major splice variants and tags SMII in all SMCs in the organism (Fig. S4). Correct targeting and expression were confirmed by genotyping (Fig. S5) and western blot analysis. Western blot analysis detected the EGFP–SMII fusion protein in heterozygous and homozygous animals, and demonstrated the expected allele-dependent shift from endogenous SMII to the higher molecular weight EGFP–SMII species (Fig. 5B). In intact arterioles in tissue explants, EGFP–SMII signal exhibited the expected circumferential/longitudinal filament architecture across the vessel wall, and optical sectioning highlighted consistent labeling across the dorsal, lumen, and ventral planes (Fig. 5C &D), supporting robust incorporation of the tagged SMII protein into native contractile structures. Finally, super-resolution SIM imaging in primary SMCs revealed SM1A head domains are about 300 nm apart flanking NM2B tails, again suggesting co-assembly of EGFP–SMII with non-muscle myosin II (Fig. 5F &G). Mice weights, blood pressures, and echocardiographs were taken to characterize cardiovascular health at 1 year of age (Fig. S6). No significant differences were detected between wildtype, heterozygote, and homozygote EGFP-SMII animals, suggesting the EGFP-SMII does not alter cardiovascular physiology. Together, these validation data establish the EGFP–SMII knock-in mouse as a physiologically relevant platform to quantify SMII assembly, exchange kinetics, and higher-order organization in primary differentiated SMCs and intact vascular tissue.

**Fig. 5.**
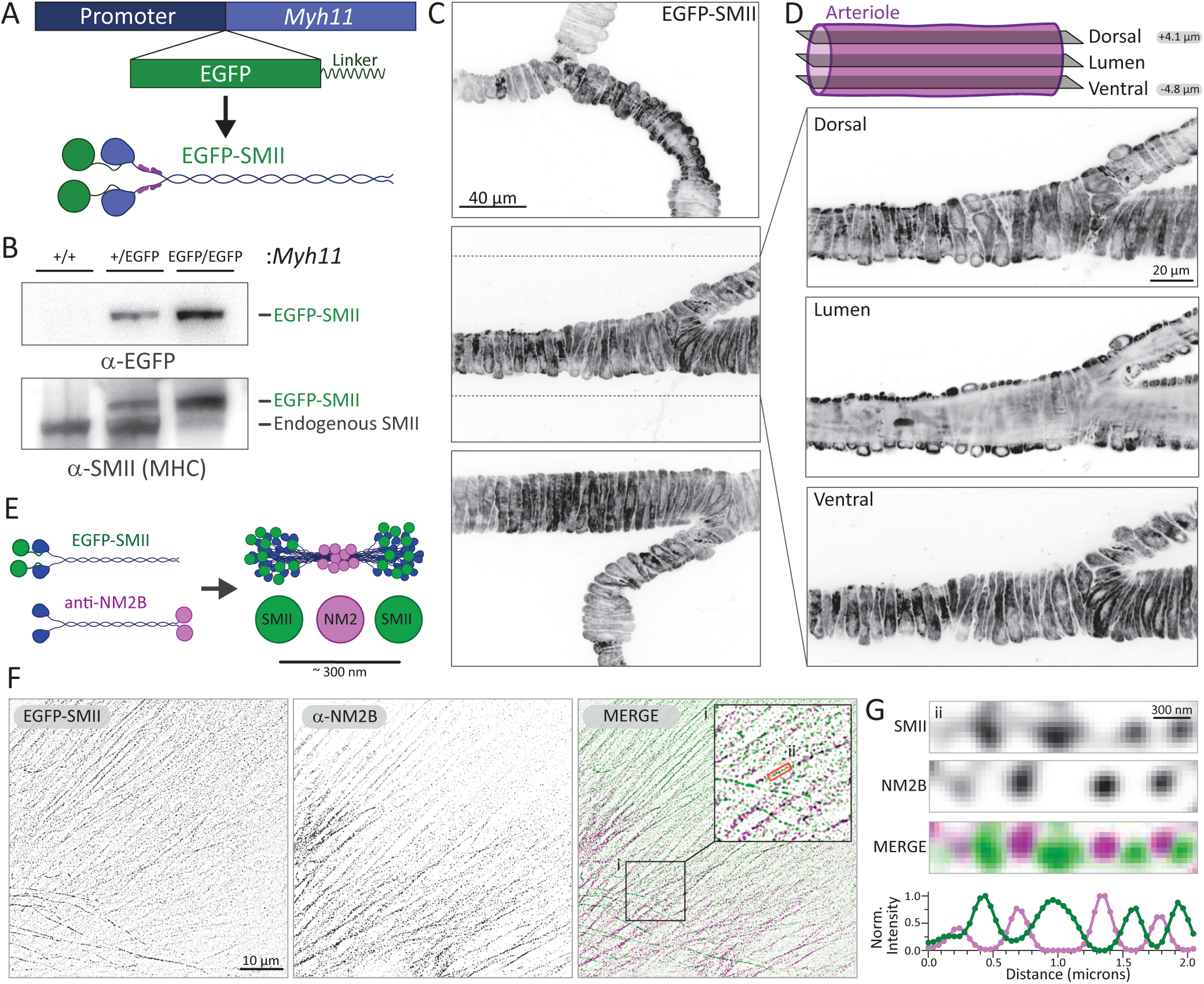
CRISPR-KI EGFP-SMII murine model reveals SMII and NMII co-assemble. **A)** Diagram of CRISPR-KI EGFP-SMII murine model. **B)** Western blot of SMC tissue from wildtype, heterozygote, and homozygote EGFP-SMII (F,M,M) mice probed with *α*-EGFP or *α*-SMII. **C)** Example images of isolated arterioles ex-vivo expressing EGFP-SMII from homozygote mice (F, M). Images are max projections of confocal stacks. **D)** Example Z-stacks were taken longitudinally to demonstrate the intact cylindrical architecture. **E)** Diagram of co-assembled EGFP-SM1A and NM2B, with the NM2B stained with a C-terminus antibody. **F)** Primary SMII-EGFP SMCs were isolated, immunostained for NMIIB, imaged with SIM, and sum projected. **G)** An individual actomyosin fiber was cropped from the cell in (F) and displayed at higher zoom. The lower plot is a line scan through the fiber normalized to maximum intensity for each channel.

### SMII filament dynamics in primary SMCs and arteriole tissue are stabilized upon transiently-induced contraction

To evaluate whether the changes in SM1A filament dynamics we observed with transiently-induced contraction in A7R5s were present in SMC physiologically, we repeated FRAP experiments on isolated primary SMCs and intact arterioles from EGFP-SMII knock-in mice (Fig. 6A). Primary SMCs were isolated from aortic tissue and cultured in vitro for up to 4 passages to limit dedifferentiation. Esophageal tissue was isolated within 2 hours of euthanasia, secured on a glass slide, and submerged in media. For primary SMCs in culture, paired control and treatment FRAP was performed in the same cell. For arterioles within esophageal tissue, we performed control FRAP on a small region of an EGFP-SMII SMC embedded in an arteriole, followed by angII treatment and a second FRAP measurement after ∼2 min (to allow agonist penetration) on a neighboring SMC within the same arteriole. This provided vessel-specific internal controls to monitor relative changes in assembly dynamics.

**Fig. 6.**
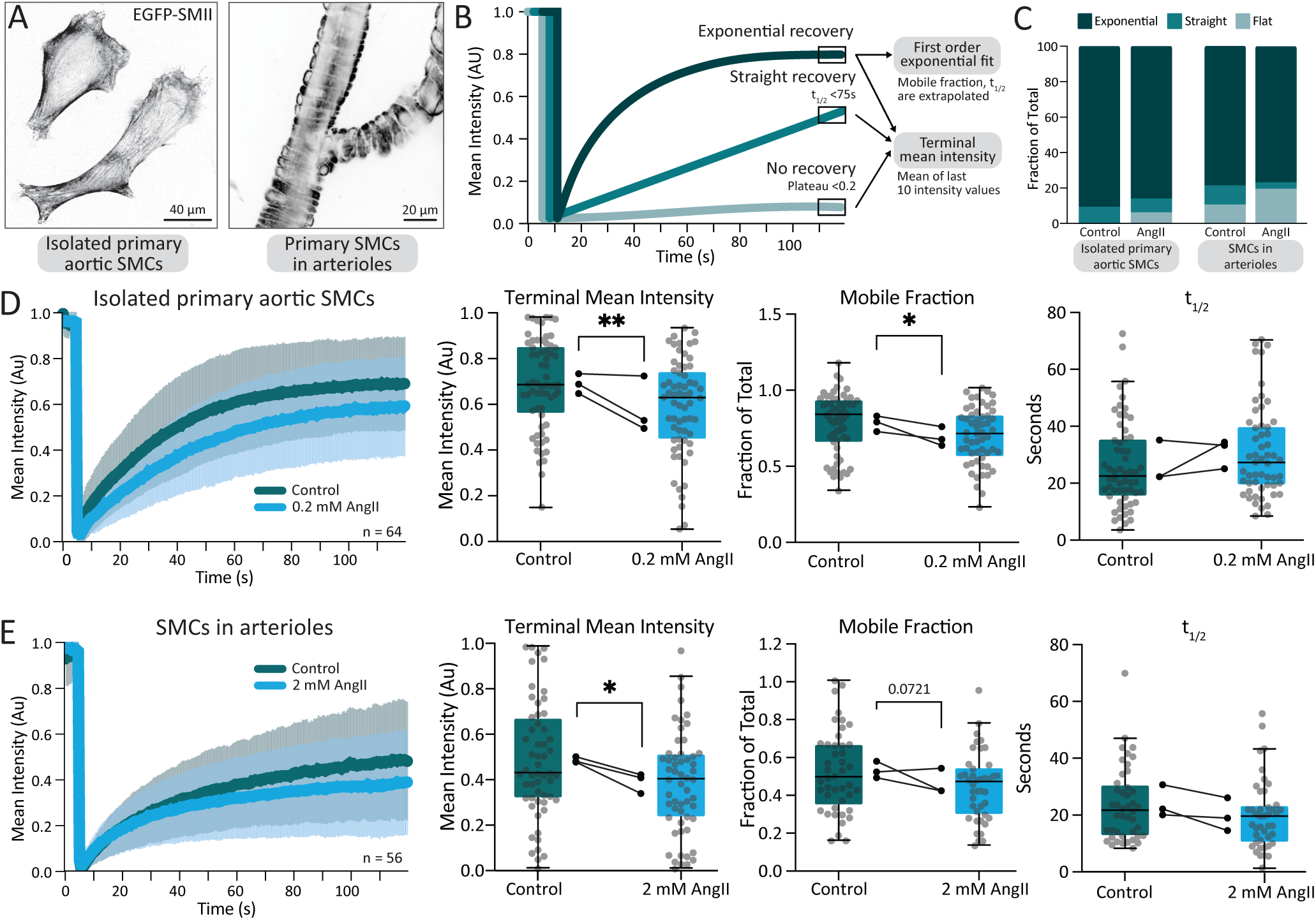
Induced contraction of primary SMCs stabilizes EGFP-SMII filaments. **A)** Example images of primary EGFP-SMII isolated vascular SMCs (left) and whole arterioles ex-vivo (right). **B)** Diagram representing the type of recovery curves we observed during FRAP experiments on primary SMCs (see also Fig. S7), and the workflow for analysis. **C)** Stacked bar graph representing the fraction of recovery types (exponential, straight, flat) we observed for each experimental group. **D)** FRAP experiments were performed on primary isolated EGFP-SMII cells. Mean recovery curves from 3 independent experiments are plotted (primary SMCs were isolated from 2 females, 1 male). Mobile fraction and t^1^⁄2 are reported with individual points in box and whiskers (median *±* quartiles). Means from each experiment are reported as dots in a spaghetti plot where pre- and post-Ang II measurements for individual experiments are connected. Paired t-tests were formed performed in control and drug treatment cells when measuring mobile fraction and t^1^⁄2. **E)** Repeat experiment on SMCs in whole arterioles ex vivo (n=3, 2F, 1 M)

Compared with A7R5 cells, FRAP recovery curves in primary SMCs in culture and in tissue explants were more heterogeneous and frequently deviated from a simple single-exponential profile. Across conditions, recovery traces fell into three reproducible classes: (1) no recovery, (2) linear recovery, and (3) exponential recovery, which we subsequently analyzed (Fig. 6B). First, recovery distributions differed across the four experimental groups (chi-square test; p=0.0098), with AngII treatment showing a higher fraction of linear/no recovery (Fig. 6C), suggestive of a shift toward a more stable SMII filament during transiently induced contraction. Next, we sought to compare all three types of recovery curves across the population with the same analysis. Because t^1^⁄2 and mobile fraction are only well-defined for exponential recoveries, we quantified the terminal mean intensity (mean of the final 10 time points) for all FRAP measurements. Terminal mean intensity decreased following angII treatment in both isolated SMCs and arterioles (Fig. 6D & E), consistent with reduced SMII exchange. Finally, we analyzed the mobile fraction and t^1^⁄2 on recovery curves that could be fit with an exponential. This once again resulted in a decreased mobile fraction, but no significant change in t^1^⁄2 (Fig. 6E), supporting activation-dependent stabilization of the SMII machinery in both isolated primary SMCs in culture and in intact arterioles.

## Discussion

Our research supports an updated model of SMII filament exchange, assembly, and contraction (Fig. 7). At steady-state, SMII is predominantly filamentous yet remains highly dynamic, exchanging with the monomeric pool, and exhibits the ability to co-assemble with NM2. Across cultured rat aorta A7R5 cells and primary SMCs and intact arterioles isolated from our EGFP-SMII KI mouse, agonist stimulation stabilizes SMII filaments, reduces SMII exchange, and increases filament assembly. Collectively, this enhanced assembly and stabilization of SMII then increases SMC contraction.

**Fig. 7.**
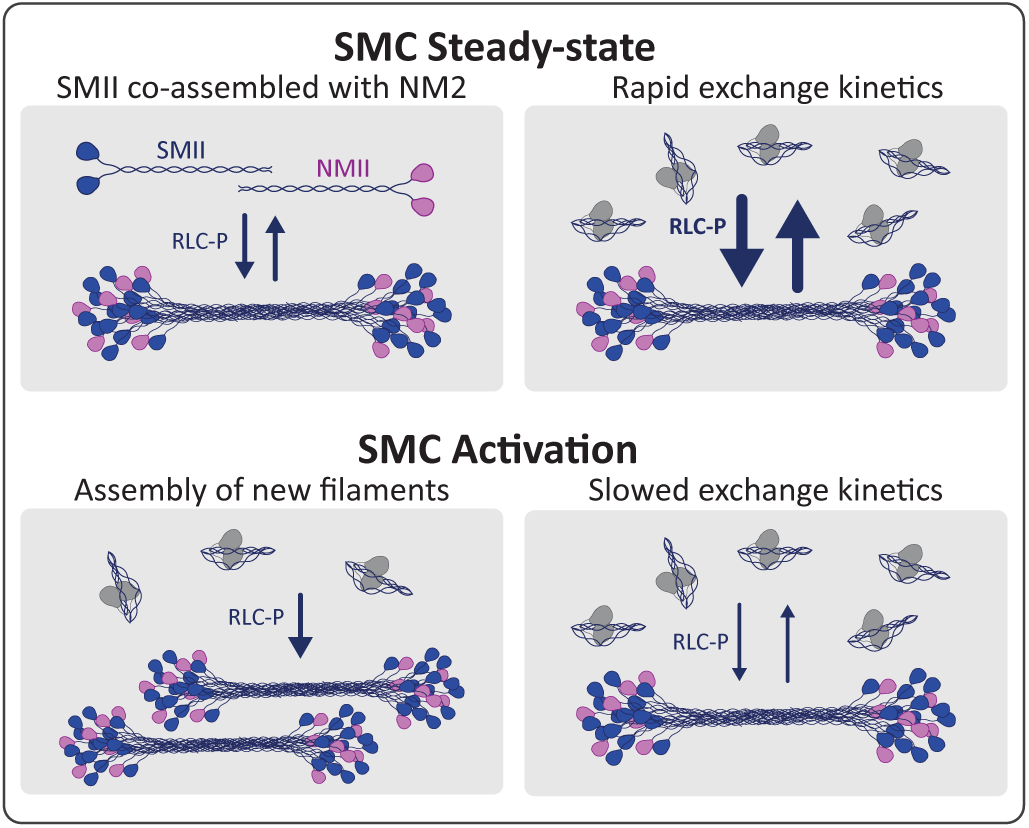
Model of SMII assembly at steady-state and upon activation. See Discussion for description.

To increase total assembly levels of myosin 2 filaments, one could increase the rate at which monomers are added (“on-rate”), or increase the dwell time in which monomers stay in the filament, or both. To increase the on-rate, the most likely scenario is that MLCK-mediated RLC phosphorylation drives the folded, autoinhibited monomer (“10S”) to open into the assembly-competent form (“6S”), which then readily incorporates into filaments (Fig. 7). This model is well-supported by the literature (18, 19, 37). To increase the dwell time, we speculate that MLCK is acting upon filamentous SMII. In theory, SMII monomers assembled into filaments could exist with 0, 1, or 2 RLCs phosphorylated, and importantly, only 1 phosphorylated RLC is required to activate the complex (38, 39). While an SMII with 0 phosphorylated RLCs might be prone to disassembly and a return to the folded 10S, early studies suggested that as much as ∼85% of SMII in smooth muscle was filamentous in relaxed smooth muscle, but only ∼5% of the RLC was phosphorylated (40). Therefore, the existence of unphosphorylated SMII monomers in filaments is likely. MLCK phosphorylation of monomers with 0 or 1 phosphorylated RLCs should extend monomer dwell time in the filament, thereby decreasing the off-rate and increasing total assembly levels. In parallel, this model of enhancing RLC phosphorylation of filamentous myosin 2 supports our FRAP data demonstrating stabilization of SMII filaments upon agonist stimulation (Fig. 4). While we find this myosin-centric model appealing, non-mutually exclusive models include stabilization by binding partners, such as caldesmon (41). Future studies that extend our dynamic imaging of SMII to include dynamic imaging of MLCK, caldesmon, and more key players should help tease apart these models.

The idea that SMII can exist as an inactive filament with relatively low RLC phosphorylation, like striated myosin 2s, but can be induced to rapidly assemble nascent filaments upon stimulation, like non-muscle myosin 2s, suggests SMII is somewhat of a hybrid of its myosin 2 paralogs. While we have a working model for how rapid assembly might occur via RLC phosphorylation of 10S monomers, described above, we have a less clear structural model of how SMII exists as an inactive filament. One possibility can be derived from studies in striated myosin 2s, where filamentous myosin 2 monomers can exist in a sequestered state, in which the motor domains dock on one another and fold back toward the proximal tail (42, 43). This interacting-heads motif (IHM) was actually first discovered in SMII and is a stabilizing feature of the folded 10S monomer (42–44). However, the IHM has not been directly observed in filamentous SMII, only in the folded monomer. Detailed analyses of the striated IHM has been enabled by the crystalline nature of striated filaments and advances in structural biology (i.e. cryo-EM) (45, 46). It is possible that the less ordered nature of SMII filaments in SMCs precludes filamentous IHM detection in cells, or that a requisite stabilizing binding partner precludes filamentous IHM detection in vitro. Whether SMII also adopts a filamentous IHM state, therefore, remains an open question.

Extracting more molecular-level assembly information from our FRAP data during induced contraction is challenged by the timescales at play. In our experiments, cytosolic calcium, SMII assembly, and traction forces all peak within 1-2 minutes of carbachol/angII-induced activation, after which they decay to steady-state levels within minutes (Fig. 3). However, plateaus in our carbachol-treated FRAP data (required to fit exponential curves) require observation for ∼5-10 minutes (Fig. 4, 6). Therefore, the SMII networks in the bleach regions are likely experiencing exchange, while simultaneously undergoing concurrent assembly and disassembly during the acquisition, complicating interpretation of a first exponential kinetic model. To address this, we quantified terminal mean intensity and curve type (exponential, linear, or minimal recovery), and we observed a consistent decrease in endpoint recovery following agonist exposure, suggesting stabilization and a reduced exchangeable pool even in cells that could not be fit exponentially. This analysis provides a conservative readout of stabilization that complements mobile fraction estimates derived from exponential curves. Future experiments with higher temporal resolution (e.g., single-molecule tracking of SMII incorporation lifetimes) should more fully elucidate changes in exchange kinetics throughout an induced contraction event.

The observation that SMII and NM2 are likely co-assembling in filaments is not unexpected, as we know that NM2 isoforms co-assemble with one another and that SMII and NM2 isoforms share high sequence and structure similarities (30, 47). While NM2 is expressed in many SMCs (24), SMII expression is dominant over NM2 (48). Therefore, the fraction of NM2 in mixed NM2:SMII filaments is likely low. Interestingly, in vitro experiments analyzing the processive movement of purified NM2 filaments demonstrated that adding a fraction (∼30%) of the NM2B isoform into filaments of the NM2A isoform was sufficient to completely render the filament NM2B-like (49). Therefore, biophysical properties of myosin 2 filaments are not linearly derived from their isoform composition, and a fraction of NM2 could meaningfully contribute to SMII filaments. Previous studies have implicated NM2 function in SMCs (50, 51). However, it is difficult to discern if any potential NM2 functionality is due to NM2 co-assembling into SMII filaments or behaving independently. Future studies selectively ablating NM2 isoforms in smooth muscle tissues or performing mixed in vitro experiments similar to previous studies with NM2 isoforms (49) to determine the biophysical properties of mixed NM2:SMII filaments should prove insightful.

Further complicating SMII and NM2 co-assembly is the presence of SMII splice variants in each SMC population (52) that could further contribute to variations in contractile properties. While in vitro studies have suggested these isoforms display unique assembly properties (53), future studies might explore exchange kinetics of SMII isoforms in cells and correlations with contractile properties of the SMCs in which the isoforms are expressed. Additional complexity could also arise from heterogeneous myosin 2 subcellular functions within SMCs, as recent evidence demonstrated unique subcellular localization for SMII and NM2 in freshly isolated primary cells (54). Finally, while the structure of NM2 bipolar filaments in cells is relatively well-established, the precise structural features of SMII filaments in intact cells and tissues (i.e. side-polar filaments) remain an active area of debate (55). Dissecting the unique and common biophysical properties of SMII and NM2 filaments in SMCs will likely yield important insights.

Smooth muscle is found in organs throughout the body, with each organ placing unique demands on the resident smooth muscle. For example, vascular smooth muscle performs tonic contraction to maintain vessel tone (56) while gastrointestinal smooth muscle performs phasic contraction to move food through the GI tract (57). Furthermore, vascular SMCs must perform different roles across the arterial tree: in large elastic arteries such as the aorta, medial SMCs contribute predominantly to wall mechanics and compliance (58, 59), whereas in small resistance arteries SMCs regulate vessel diameter, peripheral resistance, and local blood flow (60). The ability to rapidly tune force production via SMII modulation could enable rapid tuning of vascular tone or blood pressure to a new setpoint, and inability to do so could initiate or potentiate pathology (61).

Finally, pathogenic and likely-pathogenic variants in *MYH11* are reported in vascular and gastrointestinal disorders, including familial aortic aneurysm and dissection (TAAD) (4, 5). However, most of these variants have not received mechanistic scrutiny. In contrast, recent advances in cardiomyopathy-driving myosin variants (primarily *MYH7*) suggest some mutations are either stabilizing or destabilizing the IHM to create hypo- or hyper-contractile systems (62). Some pathologies driven by variants in the NM2A heavy chain, collectively termed *MYH9*-Related Disease, are found along the coiled-coil tail and are thought to disrupt filament assembly and dynamics (63, 64). We speculate that many of the reported SMII variants alter autoinhibitory state stability (10S/IHM) and/or filament assembly, thereby shifting the balance of filament availability, stabilization, and activation. Recent in vivo work further supports the idea that *MYH11* variation can produce distinct vascular phenotypes, including enhanced aortic growth under biomechanical stress, although the underlying SMII filament behavior and function remain undefined (65). As both hyper- and hypo-contractile SMCs could alter tissue compliance or tunability unfavorably, increasing or decreasing overall SMII assembly and activity could drive pathology. Identifying how these variants impact SMII dynamics could therefore inform our understanding of the pathologies they are tied to.

Overall, our studies provide foundational insight into SMC physiology by establishing that SMII filaments are both predominantly filament-associated and dynamically exchanging at baseline, and that agonist stimulation induces filament assembly and stabilization, alongside motor activation to produce force. By extending these measurements from an immortalized cell culture model to primary SMCs and intact arterioles at endogenous expression, our studies open the door into elucidating precise and unexplored mechanisms (e.g. heavy chain phosphorylation, differential MLCK signaling) for both normal and patho-physiology.

## Materials and Methods

### Mammalian Expression Plasmids

To make pEGFP-SM1A, a single gBlock was purchased from IDT that contained the 5’ 420 basepairs of human SM1A, including the naturally occurring Sal1 restriction site, a short 5 basepair linker to facilitate restriction enzyme digestion, and the terminal 3’ 616 base pairs, including the naturally occurring Blp1 restriction site and a terminal stop codon. This dsDNA was inserted into pEGFP-C1 after digestion with Bgl2 and Kpn1 restriction enzymes. The internal coding sequence of *MYH11*, not included in the gBlock, was digested from full-length cDNA (MHS6278-202857900; Horizon Discovery) with Sal1 and Blp1 (4911 basepairs) and ligated into the pEGFP-C1-SM1A intermediate following digestion with Sal1 and Blp1. To make pLVX-GCaMP7s, jGCaMP7s was PCR amplified from pGP-CMV-jGCaMP7s (Addgene #104463). Following gel extraction, this dsDNA was inserted into the pLVX backbone with a CMV promoter using Gibson Assembly. GCaMP7s subject to PCR was sequence validated. NM2A-mApple contains an mApple fluorophore on the C-terminus of NM2A, and was described previously (30).

### Cell Culture, Transfections, Transductions

Rat aortic smooth muscle cell line, A7R5 cells, were obtained from ATCC and cultured in DMEM (MT10013CV, Corning) supplemented with 10% fetal bovine serum (MT35-010-CV, Corning) and 1% antibiotic–antimycotic solution (MT30004CI, Corning). At 24 hours prior to each experiment in Figures 1-3, A7R5 cells were transfected with 2 ug total pDNA using the lipid-based transfection system LipoD293 DNA transfection reagent (#SL100668; SignaGen). For FRAP experiments in Figure 4, A7R5 cells were transduced with SM1A-adenovirus 48 hours prior at 1:2000. The A7R5 cells used for traction force microscopy were treated with lentivirus expressing GCaMP7s. A GCaMP7s positive population was obtained using fluorescence-activated cell sorting. Lentiviral production was performed in HEK-293-FT cells using psPAX.2 and pMD2.G with LipoD293 DNA transfection reagent. Lentiviral-containing media was collected at 48 and 72 hrs, filtered, and directly used for transduction of A7R5 cells. Plasmid psPAX2 and pMD2.G were gifts from Didier Trono (Addgene plasmids #12260, #12259). A7R5 cells were induced towards the contractile phenotype using serum starvation, whereby plated cells previously grown in full (10% serum) were given either 0% serum, 2.5% serum, or the control 10% serum for 24 hours (66)).

### Microscopy

Imaging was performed on the following systems. Figures 1, 2C-D, and 3A-C were acquired on a Zeiss LSM 880 Airyscan with a 63X 1.4 NA Plan-Apochromat objective in Airyscan Fast mode at 1x Nyquist sampling for optimal confocal resolution at 1 Hz for up to 1200 seconds. For FRAP experiments a circular region (5 *μ*m diameter) was bleached using 405 nm, 458 nm, and 488 nm lasers at 100% laser power. Figures 4, 5C-D, and 6 were acquired on Yokogawa W1 Spinning Disk Confocal Microscope attached to a Zeiss Axiovert7 stand with Phasor spatial illumination system (Intelligent Imaging Innovations) with a 63X 1.4 NA Plan-Apochromat objective. For FRAP experiments a 4 *μ*m circular region was illuminated at 100% laser power using the 405 nm laser for 500 ms (cells) or 1500 ms (tissue). Figures 2A-B and 5F-G were acquired on a Zeiss SIM5 with a 63x 1.4 NA Plan-Apochromat objective. Scan Mode was FastFrame with 1.0X zoom and reconstruction was performed with SIM^2^. All live-imaging was performed in environmental chambers to maintain temperature at 37*^◦^*C and 5% CO_2_, while Definite Focus (Zeiss) was used to maintain the focal plane.

### Single cell assembly assay

The single-cell assembly assay was performed on A7R5 cells transiently overexpressing EGFP-SM1A. Cells were plated in a 96-well coverglass-bottom dish (#655891, Greiner). Total EGFP-SM1A signal was collected for 16 positions per well. If applicable, small molecules were added and cells incubated for the indicated time. Then, triton buffer (5 uM PEG800, 100 mM PIPES, 150 mM NaCl, 1 mM EGTA, 1 mM MgCl2, 0.5% triton) was added to permeabilize the cells and release soluble EGFP-SM1A. The exact same positions were re-imaged to collect the triton-insoluble EGFP-SM1A signal. Individual cell masks were manually identified using FIJI. Local background fluorescence and mean fluorescence intensity was measured for each cell before and after permeabilization. Fraction assembled was then calculated as background subtracted signal post-permeabilization relative to background subtracted signal pre-permeabilization.

### FRAP

Fluorescence recovery after photobleaching (FRAP), was performed either on a Zeiss LSM 880 Airyscan (Figures 1-2), or a Yokogawa W1 Spinning Disk Confocal Microscope (Figures 4, 6). See the microscopy section for additional details. For carbachol experiments, a FRAP experiment was first performed in one region, carbachol was added, and then a FRAP experiment was performed in a new unique region. The delay between carbachol addition and initiation of the second FRAP time-course was 30-60 seconds. FRAP analysis was performed using a script written in Python similar to previous protocols (67). Three ROIs were measured for each experiment - a bleach region, a control region within the cell, and a background region outside of the cell. Intensity was monitored over time in each region. Recovery within the bleach region was obtained after normalizing to the control region and subtracting the background region. Recovery curves were fit to a single exponential *I*(*t*)= *A* ∗ (1 − *e^−kt^*), to determine the mobile fraction *A*, and *k_obs_*. The t^1^⁄2 was subsequently calculated as ln 2*/k_obs_*.

### Immunocytochemistry

Cells were fixed using a 4% paraformaldehyde solution (15710; Electron Microscopy Sciences) in cytoskeletal buffer (CB) (0.1M MES, 0.03M MgCl2, 1.38M KCl, 0.02M EGTA in 1L H2O) for 15 min at 37°C. The PFA solution was removed, cells were rinsed with 1xPBS, permeabilized with a solution of 0.5% Triton X-100 (Fisher BioReagents) in CB for 15 min, rinsed in 1xPBS, and blocked with a solution of 10% goat serum in 1xPBS for 30 min. Fixed samples were incubated with primary antibodies at 4°C overnight in 5% goat serum, washed in 0.05% Triton X-100 in 1xPBS for 3×10 min with gentle rocking, incubated in secondary antibodies diluted in blocking solution for 1 hr at room temperature, and washed in 0.05% triton in PBS 3×10 min before imaging or mounting. Cells were mounted using ProLong Glass antifade mountant (P36980; ThermoFisher). Mounted samples were allowed to cure 18 hrs in a dark drawer at room temperature prior to imaging. A7R5 cells were stained expressing EGFP-SM1A and/or NM2-mApple and 405 phalloidin (A30104, ThermoFisher). Primary EGFP-SMII SMCs were stained for NMIIB (# 230823, AbCam) and phalloidin.

### Western blot

Cells were scrape collected or pelleted and resuspended in 2x sample buffer, boiled 5 min, sonicated briefly, and used for analysis or stored at -20*^◦^*C. Tissues were weighed out to ∼30-50 mg, cut up into small pieces, resuspended in 2x sample buffer, boiled 10 min, sonicated briefly, and used for analysis or stored at -20*^◦^*C. Proteins were separated on 4-20% gel, unless separation of EGFP-SM1A and SMII was required, in which case a 7.5% gel (cells) or 3-8% tris-acetate gel (tissues) was used. Protein was transferred to a PVDF membrane using eBlot L1 (Genscript). Blocking was performed with 5% milk in PBST (PBS + 0.1% TritonX-100) and primary incubations were done overnight at 5*^◦^*C. Primary antibodies were all used at 1:2,000 in blocking buffer (SMII, Abclonal #A4064; SMA Abclonal #A17910; EGFP, Santa Cruz #SC-9996)

### Traction force microscopy

Traction force microscopy was performed as described previously (68, 69) on the Spinning Disk - see microscopy section. Coverslips were prepared by incubating with a 2% solution of 3-aminopropyltrimethyoxysilane (313255000, Acros Organics) diluted in isopropanol, followed by fixation in 1% glutaraldehyde (16360, Electron Microscopy Sciences) in ddH20. Polyacrylamide gels (shear modulus: 16 kPa—final concentrations of 12% acrylamide [1610140, Bio-Rad] and 0.15% bis-acrylamide [1610142, Bio-Rad]) were embedded with 0.04 *μ*m fluorescent microspheres (F8789, Invitrogen) and polymerized on activated glass coverslips for 1 hrat room temperature. After polymerization, gels were rehydrated overnight and coupled to human plasma fibronectin (FC010, Millipore) for 1 hrat room temperature using the photoactivatable cross-linker Sulfo-Sanpah (22589, Pierce Scientific). Following fibronectin crosslinking, cells were plated on the gels and allowed to adhere overnight. The next day, images were taken of both the cells and the underlying fluorescent beads. For kinetic analyses, images were collected every 30 s for 30 min. The first 10 min captured the steady-state, then 10 *μ*M carbachol was added and the cells were imaged for another 20 min. Immediately after imaging, cells were removed from the gel using 0.05% SDS and a reference image of the fluorescent beads in the unstrained gel was taken.

Analysis of traction forces was performed using code written in Python according to previously described approaches (available at https://github.com/OakesLab/TFM). Prior to processing, images were flatfield corrected and aligned to the reference bead image. Displacements in the beads were calculated using an optical flow algorithm in OpenCV (Open Source Computer Vision Library, https://github/itseez/opencv) with a window size of 8 pixels. Traction stresses were calculated using the FTTC approach, with a regularization parameter of 6.1 × 10*^−^*^4^. The strain energy was calculated by summing one-half the product of the strain and traction vectors in the region under the cell (70) and normalized by the cell area as measured using the GCaMP7s image of the cell.

### General animal maintenance and welfare

All animal procedures were approved by the Institutional Animal Care and Use Committee (IACUC) at Loyola University Chicago and conducted in accordance with the NIH Guide for the Care and Use of Laboratory Animals and institutional guidelines. Mice were housed in individually ventilated cages under standard conditions (12-hour light/dark cycle), temperature, humidity, and with access to standard chow and water. Health status was monitored daily by trained animal care staff, and all procedures were performed with efforts to minimize the pain and distress of animals. **Generation of EGFP-SMII knock-in mice, breeding, and genotyping.** Initial CRISPR/Cas9 knock-in was performed at the University Chicago Transgenic Mouse Facility. C57BL/6J females were superovulated using standard PMSG/HCG procedure. 1-cell fertilized eggs were isolated on the day of injections. CRISPR/Cas9 components were delivered via pronuclear microinjection into 1-cell fertilized eggs. The ribonucleoprotein (RNP) complex was prepared by annealing synthetic crRNA and tracrRNA (5’-CAGGGGACCACCAGACATCA-3’ for *Myh11*; IDT) at a 1:1 molar ratio to form the sgRNA complex, which was subsequently incubated with Alt-R S.p. Cas9 nuclease (IDT) at room temperature for 20 min to ensure stable complex formation. The final injection mixture consisted of 25 ng/μL sgRNA, 100 ng/μL Cas9 nuclease, and 12.5 ng/μL donor plasmid DNA in cell culture grade ddH2O. The donor plasmid (pUC57-HA-EGFP-MYH11-Ms) consisted of a 705 bp 5’ HDR of *Myh11* genomic sequence immediately upstream of the endogenous start codon (5’-TATGTGAGTA…ACCAGACATC-3’), an HA tag, a short 7 amino acid GS-rich linker, mEGFP, a linker, an 11 amino acid linker, and 714 bp 3’ HDR of *Myh11* genomic sequence (5’-ATGGCGCAGA…CTCTGCCATG-3’). Following microinjection into the male pronucleus, surviving embryos were cultured briefly and then surgically transferred into the oviducts of pseudopregnant recipient females to allow for gestation and birth of founder mice. Founder mice were screened for EGFP insertion and bred out to screen for germline transmission.

The EGFP-SMII knock-in line was expanded by breeding the founders with wild-type C57BL/6J mice, and subsequently bred to homozygosity. To reduce background variability, the line was backcrossed for three generations prior to the establishment of homozygote animals. Genotyping was performed using ear-clip tissue obtained at weaning. Ear clips were digested using an acid-based lysis/digestion method (25 mM NaOH, 0.2 mM EDTA pH = 12 at 95C for 30 min, neutralized with 40 mM Tris-HCl pH=5 on ice) followed by PCR with primers flanking the integration site (Fig. S5). PCR products were analyzed by agarose gel electrophoresis to identify wild-type, heterozygous, and homozygous genotypes.

### Body weight, blood pressure, and echocardiography

To assess baseline cardiovascular phenotype in our novel murine model, measurements were performed at 1 year of age to increase sensitivity for detecting genotype-associated differences in blood pressure and cardiac function. Body weights were recorded. Noninvasive systolic blood pressure was measured using a tail-cuff system (Kent Scientific CODA Monitor). Mice were acclimated to restraint and tail-cuff measurements for 2-3 consecutive days prior to data acquisition, and systolic blood pressure was calculated from the average of 15 accepted cycles per session. Transthoracic echocardiography was performed using a high-frequency Vevo 3100 ultrasound system. Mice were anesthetized with inhaled isoflurane and maintained on a heated platform with continuous monitoring of temperature, heart rate, and respiration. Left ventricular systolic function (including ejection fraction) and mitral valve inflow parameters were quantified from standard parasternal long- and short-axis views and Doppler recordings (M-mode and B-mode). Analyses were done using Vevo LAB ultrasound analysis software.

### Isolation and culture of primary EGFP-SMII vascular SMCs from aorta

To isolate primary EGFP-SMII SMCs, homozygous EGFP-SMII mice (12-16 weeks of age, male and female), were administered deep anesthesia followed by exsanguination. Briefly, the abdominal cavity was opened, the inferior vena cava was transected, 1xPBS was perfused via the left ventricle until effluent cleared. The thoracic aorta was dissected under a dissecting microscope, and surrounding adipose and connective tissue were removed. The aorta was minced into small fragments and incubated in collagenase solution (collagenase type I, 10 mg/mL in PBS; Sigma-Aldrich, C0130) for 2 hours at 37*^◦^*C. Following digestion, tissue was pelleted by centrifugation (400 rcf, 4 min), resuspended in vascular SMC growth medium (C-22062, PromoCell), and passed through a 70 *μ*m cell strainer (352350, Falcon). Cells were plated in 35-mm dishes and maintained at 37*^◦^*C in 5% CO_2_. Medium was changed after 48 hours; cells were used for experiments after 72 hours and/or passaged as needed. To limit phenotypic drift, primary SMCs were discarded after passage 4.

### Ex vivo arteriole preparation and imaging

For ex vivo arteriole imaging, the esophagus was carefully dissected from homozygous EGFP-SMII mice (12-16 weeks of age, male and female), opened longitudinally, and splayed flat on a glass coverslip using a tissue harp in media (641419, Warner Instruments). Arterioles within the esophageal tissue were identified by confocal microscopy based on vessel morphology and EGFP-SMII signal. Tissues were maintained at 37*^◦^*C, 5% CO_2_ in the microscope humidity chamber, and imaged within 3 hours of euthanasia to minimize time-dependent loss of tissue viability. Bladder, colon, and uterus were also dissected from animals and flash frozen or processed for Western blots (as described earlier).

**Stats.** Statistics were performed using the GraphPad Prism. Asterisks are used to convey statistical significance are as follows: * for p < 0.05, ** for p < 0.01, *** for p < 0.001, and **** for p < 0.0001.

## Acknowledgements

This work was supported in part by National Institutes of Health grants R35-GM138183 to J.R.B, R01-GM148644 to P.W.O., F30-HL178288 to SKD, LUC Transformative Research Pilot Grant to J.R.B and P.W.O, and shared instrument grants to LUC for Zeiss SIM5 (S10OD034431) and transthoracic echocardiography equipment (S10OD028449). We thank the Transgenic Animal Facility at University of Chicago and Loyola University Chicago Imaging Facility for core support.

## Author Contributions

MAB, SKD, PWO, and JRB developed the project. SKD, MAB, BS, HW, MEU, SS, HP, ERL, and JRB performed experiments. SKD, MAB, BS, HW, AZ, JAK, and JRB generated critical reagents. SKD, MAB, BS, PWO, and JRB analyzed the data. SKD, MAB, PWO and JRB wrote the initial manuscript. All authors edited and approved the final text.

## Supplementary Information

**Fig. S1.**
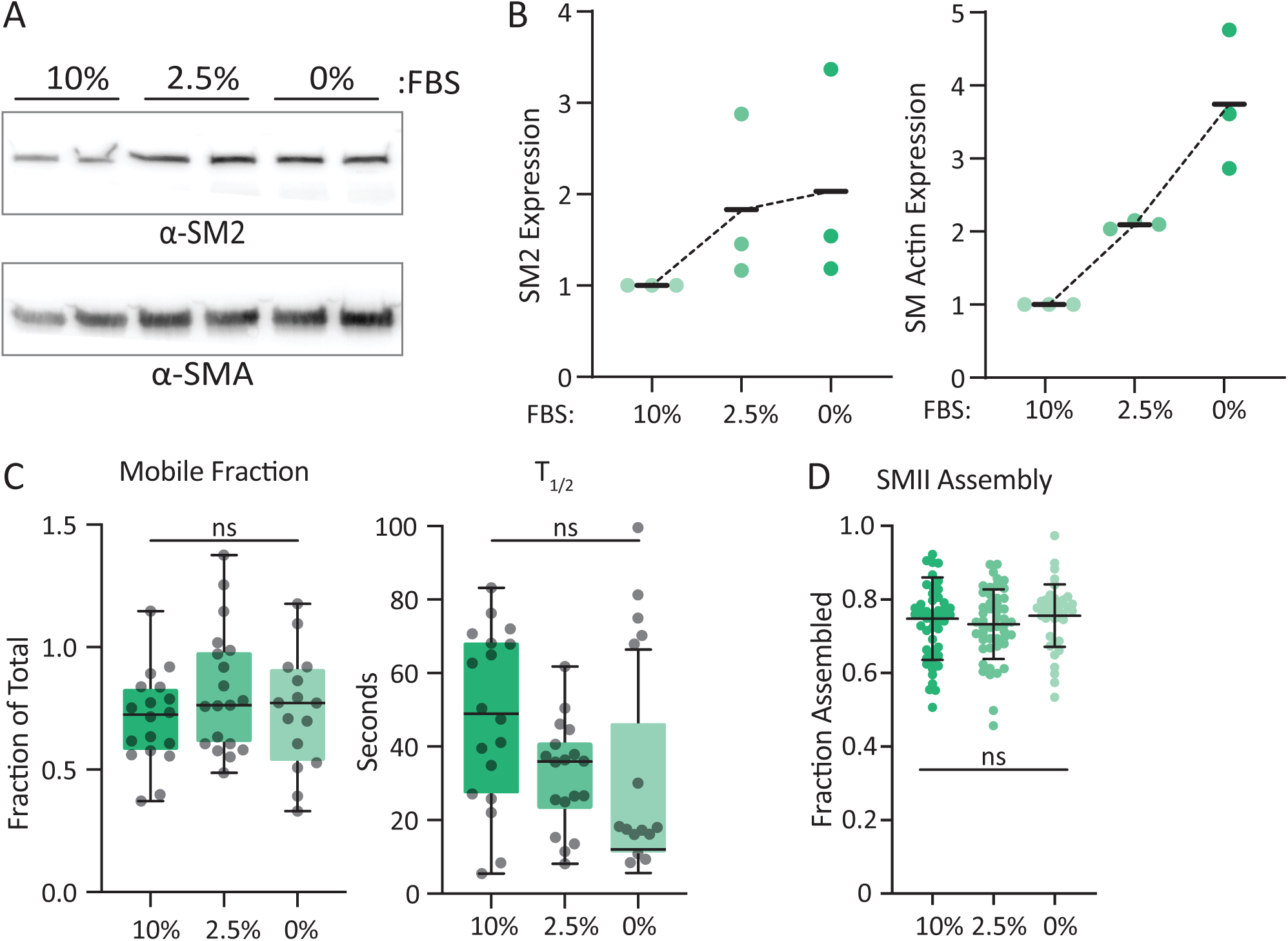
Differentiation to a more contractile phenotype does not change filament assembly or dynamics. **A and B)** A7R5 cells cultured in media with 10%, 2.5%, or 0% FBS in duplicate were subject to western blot analysis with *α*-SMII (top blot) and *α*-smooth muscle actin (bottom blot). Cumulative data plotted in B as mean *±* SD from 6 samples per condition from 3 independent experiments. **C)** FRAP was performed on A7R5 cells expressing EGFP-SM1A cultured in media containing the indicated levels of FBS. Mobile fraction and plotted as the mean of 14-18 cells per condition over 3 independent experiments. **D)** Single cell assembly assay was performed on A7R5 cells expressing EGFP-SM1A cultured in media containing the indicated levels of FBS. Data plotted from 3 independent experiments. One way ANOVA was performed between groups and showed no difference.

**Fig. S2.**
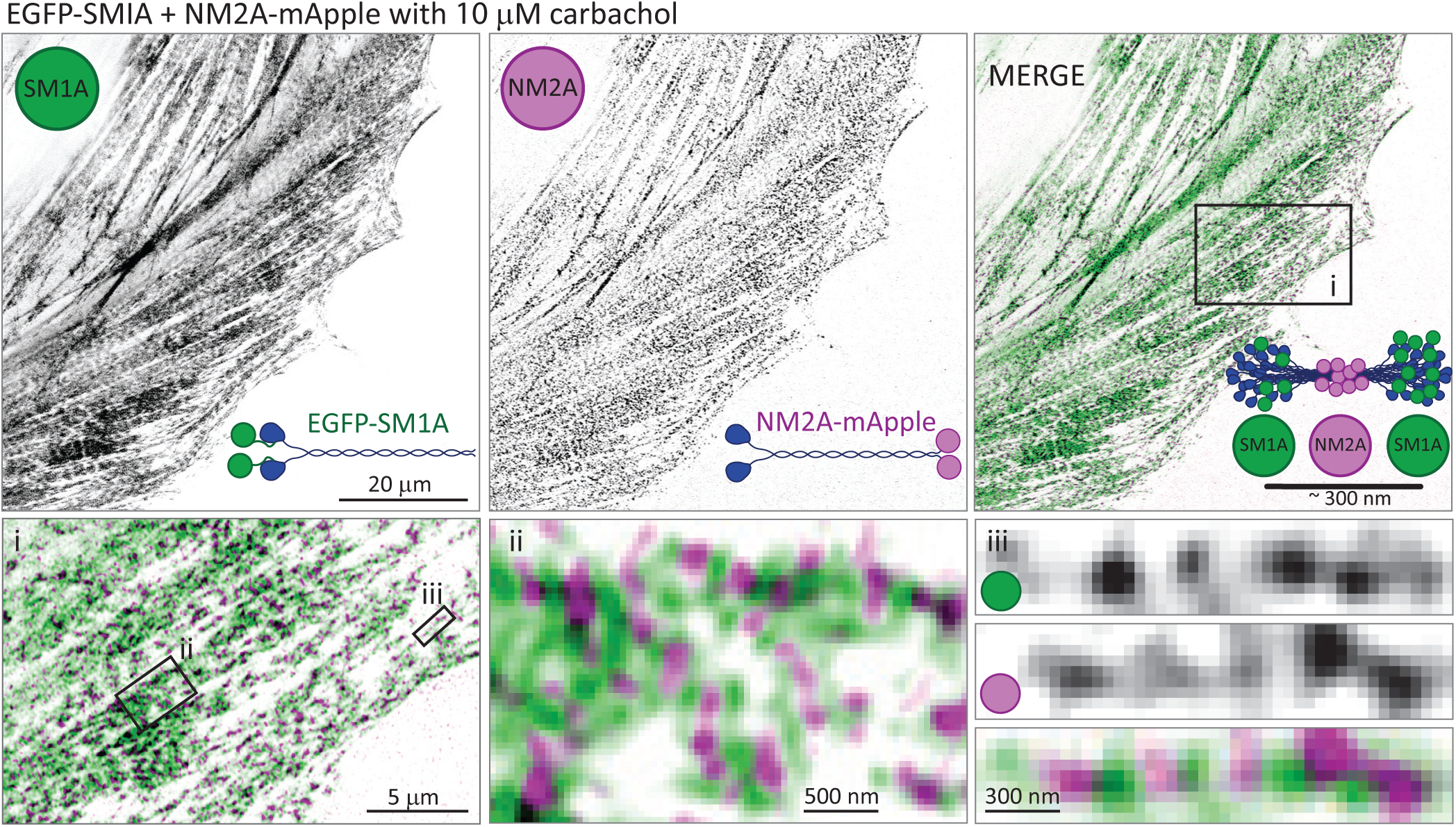
SMIA and NM2A display co-assembly upon carbachol stimulation. A7R5 cells co-transfected with EGFP-SM1A and NM2A-mApple were treated with 10 uM carbachol and fixed after 1 min. Samples were imaged on a Zeiss LSM 880 Airyscan and processed with joint-deconvolution. Single confocal slices are displayed of a representative cell. Individual channels are displayed in inverted grayscale, and merge is shown in the right panel. Diagrams of EGFP-SM1A, NM2A-mApple, and co-assembled filaments are displayed in the bottom right corners. Zoom insets are labeled ’i’ through ’iii’ and correspond to subsequent images.

**Fig. S3.**
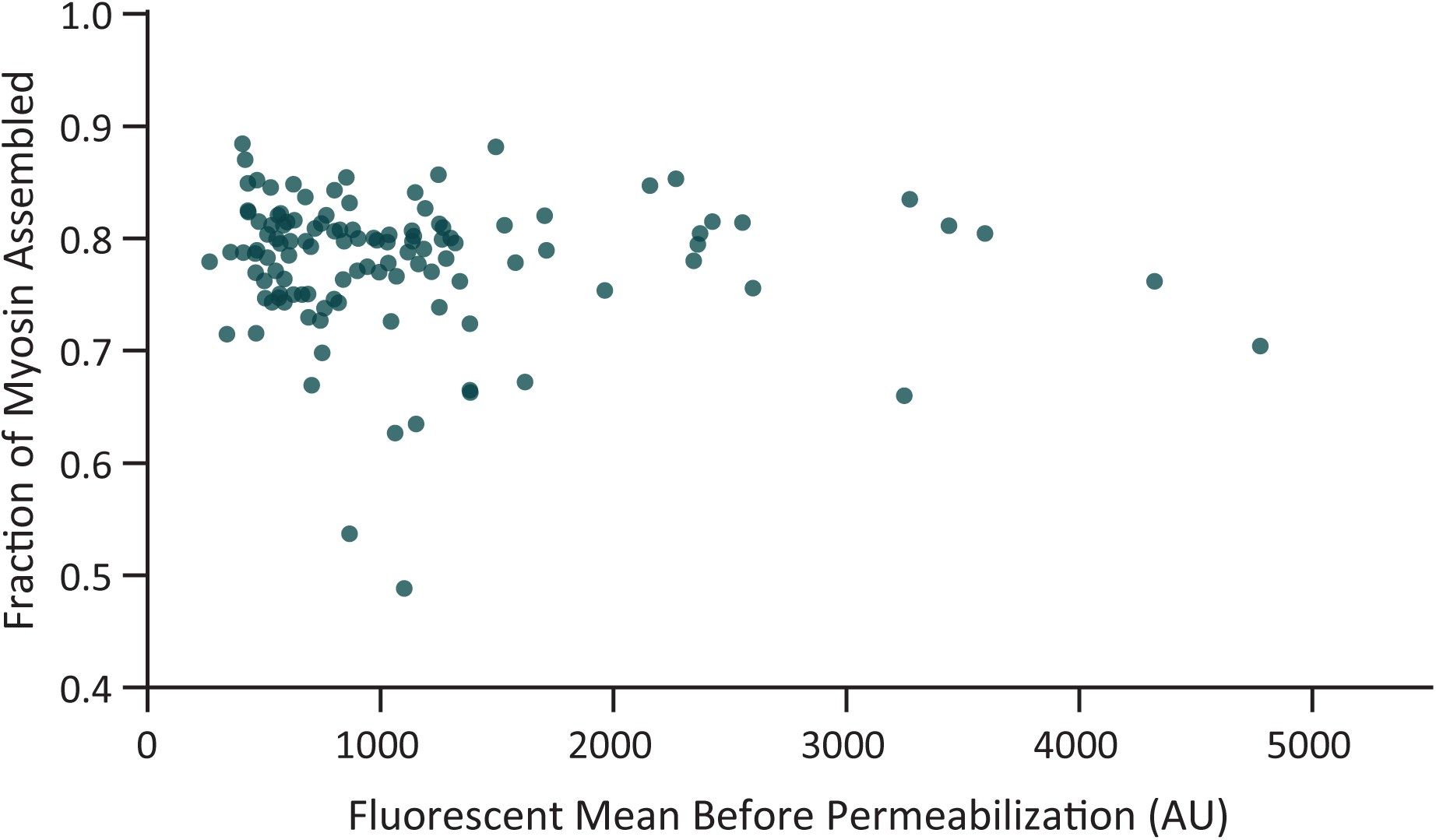
Exogenous SM1A expression does not alter assembly levels. Mean intensity of EGFP-SM1A expression prior to permeabilization x-axis plotted versus the fraction of EGFP-SM1A assembled (y-axis) demonstrates no apparent correlation between expression and assembly. This suggests the level of exogenous expression does not significantly alter the fraction of assembled filaments.

**Fig. S4.**
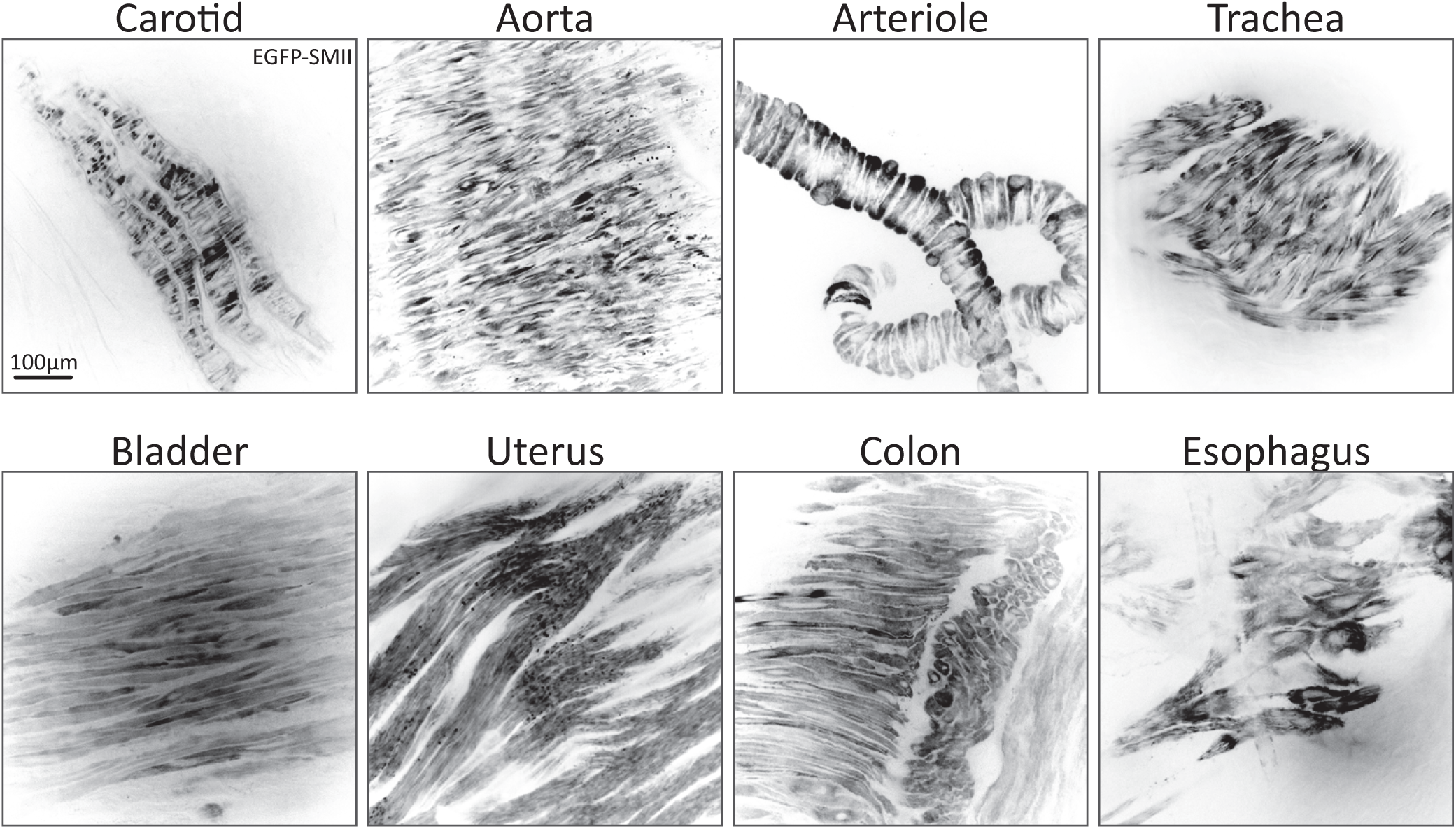
EGFP-SMII is expressed throughout the organism. Tissue explants from EGFP-SMII mice were imaged shortly after isolation (2-3 hrs). As expected, EGFP-SMII signal is apparent in all isolated smooth muscle, albeit with notably different organization. Scale bar in top left image applies throughout.

**Fig. S5.**
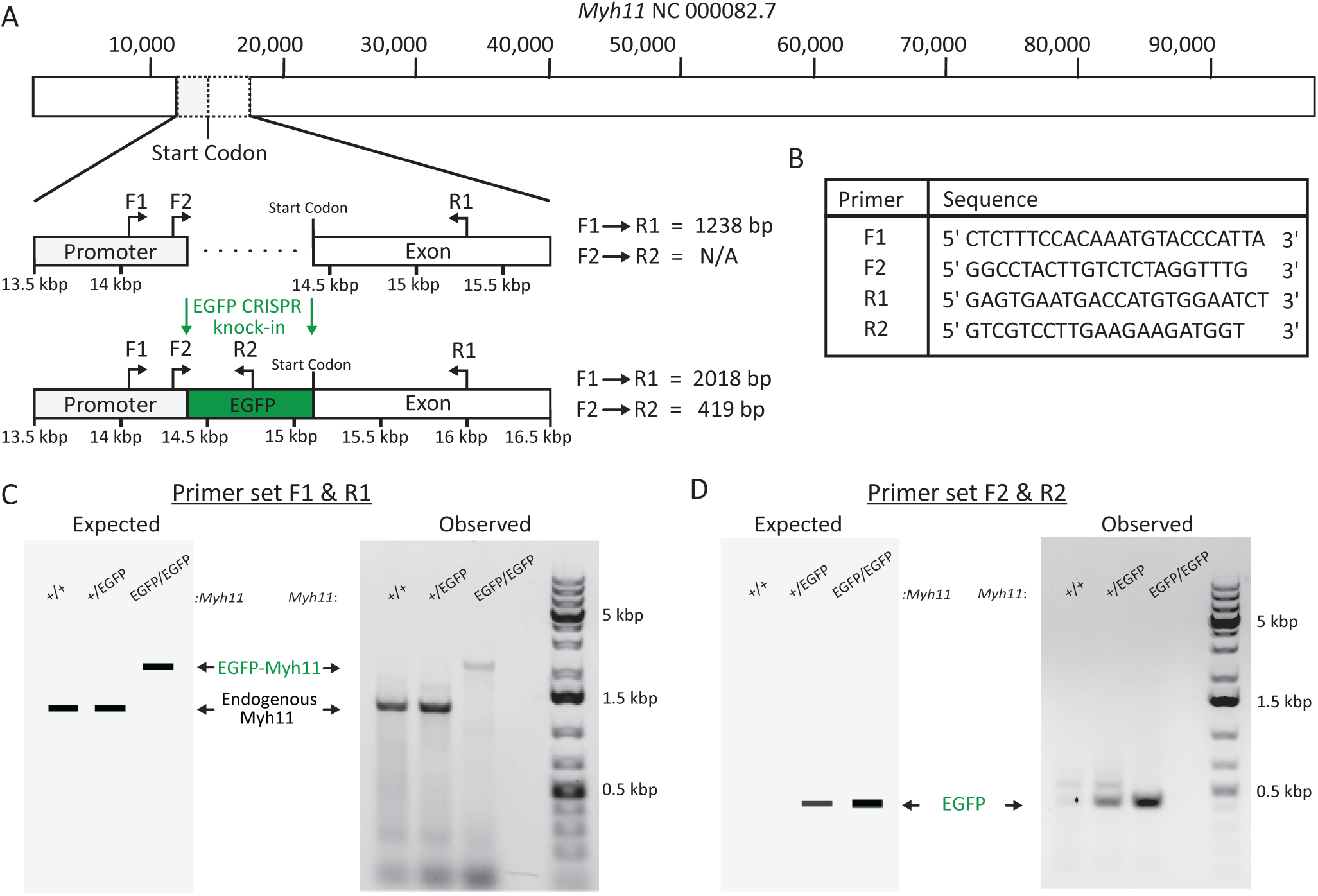
CRISPR-KI SMII-EGFP murine model genotyping. **A & B)** Diagram of genomic region of *Myh11* where EGFP was introduced via CRISPR/Cas9. PCR primer locations in (A) correspond to sequences in (B). Expected PCR product sizes for wild-type and EGFP knock-in for F1-R1 and F2-R2 primer sets are indicated. **C & D)** Expected and observed PCR products for F1-R1 and F2-R2 primer sets. Notably in (C), we continuously struggled to identify primers and PCR conditions to faithfully produce both a wild-type band and EGFP knock-in band for heterozygotes. We attribute this to the smaller wild-type PCR product preferentially amplifying compared to the larger EGFP knock-in product. Therefore, we used the second primer set (F2-R2 in D) to complement the first and validate the presence of the EGFP.

**Fig. S6.**
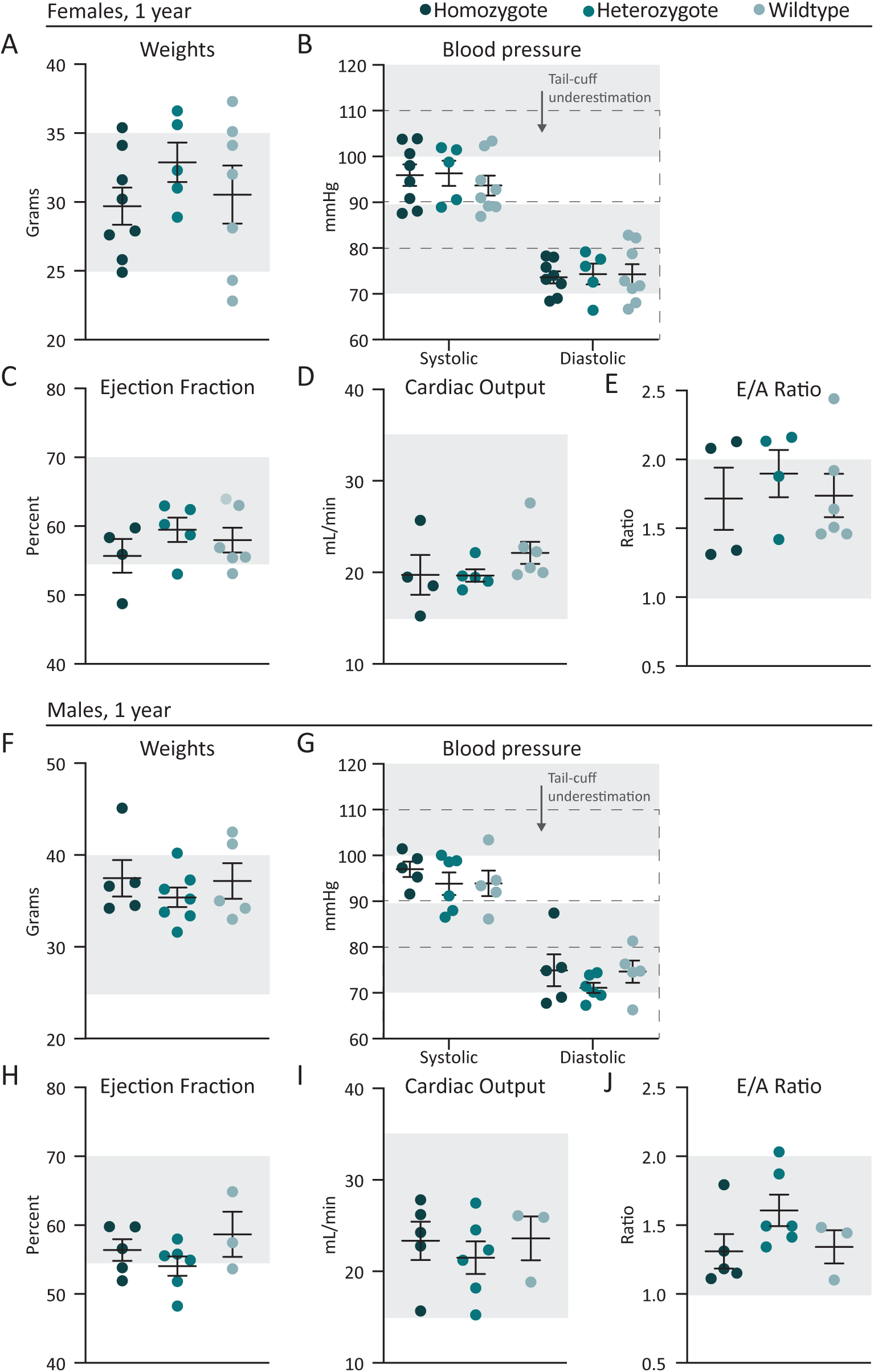
EGFP-SMII model cardiovascular measurements. Wild-type, heterogzygotic (+/EGFP-Myh11), and homozygotic KI (EGFP-Myh11/EGFP-Myh11) females (A-E) and males (F-J) were analyzed with the indicated physiological measurements. Grey regions indicate standard ranges according to the literature. We note that tail-cuff blood measurements are known to underestimate blood pressure by about 10 mmHg (indicated by lower dotted boxes in B and G). Dots represent individual animals with mean +/-SD indicated.

**Fig. S7.**
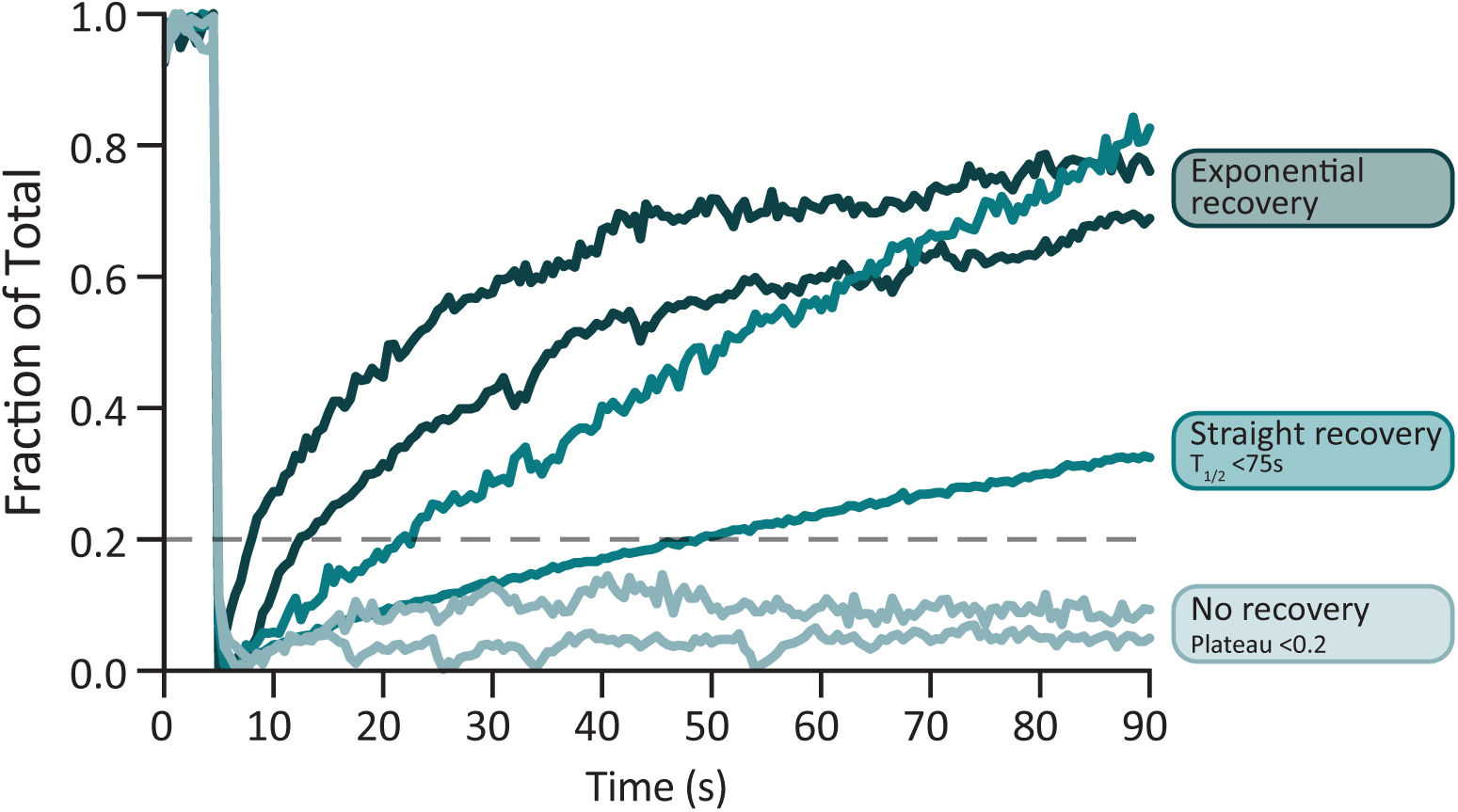
Sample FRAP recovery types. Plotted are 6 individual recovery curves (2 examples of each, as indicated by color) from the control arteriole FRAP experiment (Fig.6), displaying the (1) flat, (2) straight, and (3) exponential recovery profiles.

